# Deciphering the Network Architecture of APOBEC3-Driven Mutagenesis in HPV-Positive Head and Neck Cancers

**DOI:** 10.64898/2026.09.17.752415

**Authors:** Jake D. Lehle, Mohadeseh Soleimanpour, Niloofar Haghjoo, Colin Rorex, Feng Li, Armando Mendez, Rachael Rodriguez, Elizabeth Sommer, Cheng-Ming Chiang, Diako Ebrahimi

## Abstract

2

APOBEC3A (A3A) and APOBEC3B (A3B) are cytosine deaminases that restrict viral infection and can also mutate the host genome. In human papillomavirus (HPV)-positive head and neck squamous cell carcinoma (HNSCC), expression of both enzymes is elevated, but bulk sequencing averages their effects across mixed cell populations. Here we profile single cells from HPV16-positive HNSCC tumors and matched normal tissue. We find that the APOBEC3 (A3) single base substitution mutational signature, SBS2, is enriched in cells expressing more *A3A* than *A3B*, whereas copy number alteration (CNA) burden is enriched in cells expressing more *A3B* than *A3A*. Tumor versus normal co-expression networks identify the A3 interactors RALY and HNRNPA2B1 as candidate A3 activators. We find that SBS2 and CNA mark the maintenance and virus assembly stages of the HPV16 life cycle, and their ratio may offer a molecular estimate of tumor age. The neoantigens from immune-visible SBS2-HIGH and immune-evasive CNA-HIGH cells identify candidates for mRNA vaccines matched to a tumor’s viral state.

**Significance:** Single-cell analysis of HPV-positive HNSCC reveals that DNA editing enzymes A3A and A3B associate with distinct HPV16 life cycle stages and immunological tumor states, enabling stage-matched neoantigen vaccine candidate identification for personalized immunotherapy.

## 4 Introduction

The human genome encodes seven APOBEC3 (apolipoprotein B mRNA editing catalytic polypeptide-like 3), also known as the A3 enzymes (A3A/B/C/D/F/G/H), which are core components of innate immunity. These proteins restrict exogenous viruses and endogenous retroelements through cytosine deamination of viral genomes and also deamination-independent mechanisms^1–3^. When dysregulated, a subset of A3 family members, particularly A3A and A3B, can redirect their mutagenic activity toward the host genome. These enzymes generate characteristic C>T and C>G mutations at TCW (W:T/A) trinucleotide contexts that define the COSMIC mutational signatures SBS2 and SBS13, respectively (RRID:SCR_002260)^4–6^. A3 mutagenesis is among the most pervasive endogenous sources of somatic mutation in cancer, and its signature is detectable in roughly half of primary and metastatic tumors^7,8^.

Enrichment is strongest in virus-associated malignancies, including cervical cancer and HPV-positive head and neck squamous cell carcinoma (HNSCC)^4^. HPV infection upregulates *A3A* and *A3B* expression as part of the cellular response^9–11^. What remains unclear is how this response turns into mutagenesis of the host genome.

Most prior work has relied on bulk analysis, which averages signals across diverse cell populations and states. This averaging confounds interpretation in three ways. First, *A3* enzymes are expressed in tumor epithelial cells but also in immune cells (*A3A* in macrophages and dendritic cells and *A3B* in plasma cells and some B cells)^12^. Associations drawn from bulk expression or mutational data therefore cannot be assigned to a specific cell type. Second, mutational signatures including SBS2/13 are not uniform across cells within a tumor, so bulk-averaged signatures obscure the range of activity present across individual cells. Third, HPV infection is itself heterogeneous. Not all cells in a lesion carry the virus, and viral gene expression and life cycle stage vary substantially from cell to cell^13,14^. *A3* induction consequently varies across cells in ways that bulk measurements cannot resolve. For these reasons, it is necessary to use single-cell analysis. Single-cell studies of HNSCC have characterized malignant cell states, viral gene expression heterogeneity, and the tumor microenvironment^15–17^, and have shown that *A3A* and *A3B* are expressed in distinct epithelial cell states in normal squamous epithelium^18^. However, it remains unclear how *A3* patterns relate to the viral life cycle stage of individual cells in HPV-positive HNSCC.

Here, we analyze a single-cell RNA-seq dataset of HPV16-positive HNSCC tumors and matched normal-adjacent tissue (GSE173468)^19^, quantifying gene expression, HPV transcription, somatic mutations, mutational signatures, and predicted neoantigens. Within the basal epithelial compartment, we resolve two tumor cell populations. One carries a high SBS2 burden and expresses more *A3A* than *A3B*. The other carries a high burden of copy number alterations (CNAs) and expresses more *A3B* than *A3A*. Separating these populations yields three findings: First, tumor-specific co-expression networks centered on *A3A* and *A3B* identify *RALY* and *HNRNPA2B1*, both known A3 interactors, as candidate protein cofactors predicted to promote A3-driven mutagenesis. Second, the two epithelial cell populations correspond to distinct stages of the HPV16 life cycle, suggesting that their ratio may provide a molecular estimate of HPV16-positive HNSCC tumor age. Third, the neoantigens predicted in each epithelial cell population define candidate targets for mRNA vaccines matched to a tumor’s viral state.

One axis unifies all three findings: the ratio of SBS2 to CNA tracks the stage of the viral life cycle. A3-driven deamination predominates during the HPV16 genome maintenance stage, when *A3A* expression is highest. Progression to the later life cycle stages requires HPV16 to engage the host double-strand break response^20^, and CNA accumulate as a consequence of that repair. Cells carrying the late-stage program retain fewer detectable SBS2 mutations. This observation is consistent with deletion of the sequence carrying SBS2 mutations, decreased expression in the transcripts in which they are detected, and with selection against the most heavily mutated cells. This axis also points toward practical use, including molecular estimation of tumor age and neoantigen vaccines matched to viral life cycle state. The need for such analysis is growing. HPV-associated oropharyngeal cancer has surpassed cervical carcinoma as the most common HPV-attributable malignancy in the United States^21^, with incidence rising among younger patients^22^.

## 5 Results

### 5.1 Single-cell analysis localizes A3-driven mutagenesis to basal epithelial cells and resolves divergent *A3A*:SBS2 and *A3B*:CNA linked programs

To resolve the cell-type-specific origins of A3-driven mutagenesis, we analyzed HPV16-positive HNSCC tumors and matched normal-adjacent tissue from 14 patients (GSE173468)^19^. The dataset comprised 155,650 cells across 44 samples, 129,828 from tumor tissue and 25,822 from normal-adjacent tissue. Cell-type annotation with popV^23^ followed by cluster-level refinement resolved 12 populations (Supp. Figs. 1-3), of which 52,126 cells (33.5%) were annotated as basal epithelial. Across the A3 family, *A3A* and *A3B* were expressed predominantly in the basal epithelial compartment (*A3A*: mean = 1.62, 25.7% of cells positive; *A3B*: mean = 1.69, 33.2% positive). Lower levels of *A3A* were also detected in macrophages (mean = 0.62, 10.7% positive) and myeloid dendritic cells (mean = 0.40, 7.3% positive) (Fig. 1, Fig. 2a-c). *A3C* and *A3G* were expressed largely in immune cells, and *A3D*, *A3F*, and *A3H* showed low, diffuse expression across multiple cell types. These data are consistent with previous work identifying *A3A* and *A3B* as the family members most strongly dysregulated in basal epithelial cells^24^. We therefore focused on *A3A* and *A3B*, which are the nuclear-localized family members with access to genomic DNA.

**Figure 1:**
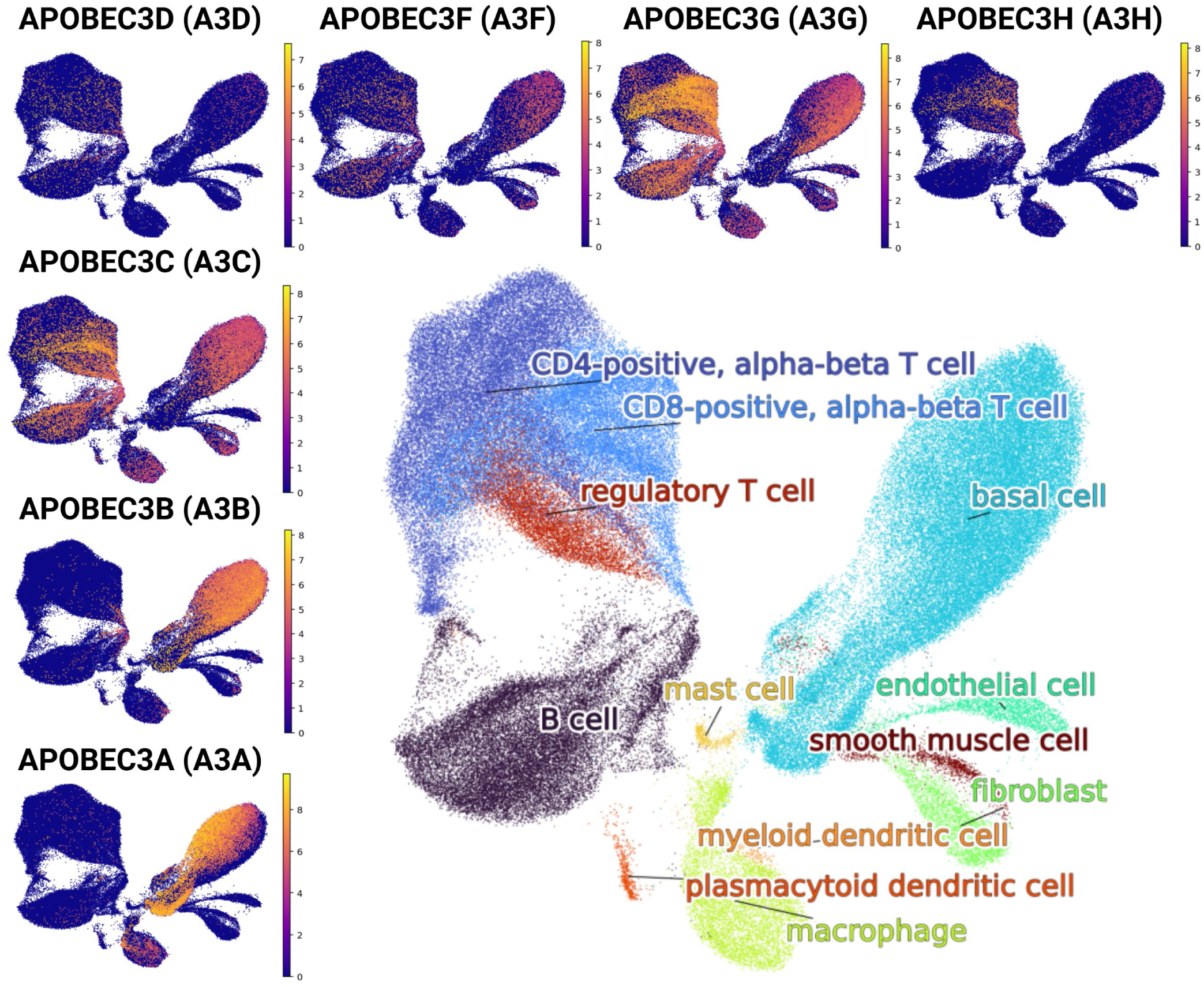
A3 family member expression across single-cell populations. UMAP embeddings colored by expression of each A3 family member: *A3A, A3B, A3C, A3D, A3F, A3G*, and *A3H*. *A3A* and *A3B* expression is concentrated primarily within the basal epithelial cell cluster and is also present in the macrophage and dendritic cell populations at lower levels, consistent with their roles as the primary drivers of SBS2 mutagenesis in identified basal cells. *A3C* expression localizes predominantly to immune cell populations. *A3G* expression is globally expressed but peaks in immune cell populations. *A3D, A3F,* and *A3H* expression is low and diffuse across multiple cell types.

**Figure 2.**
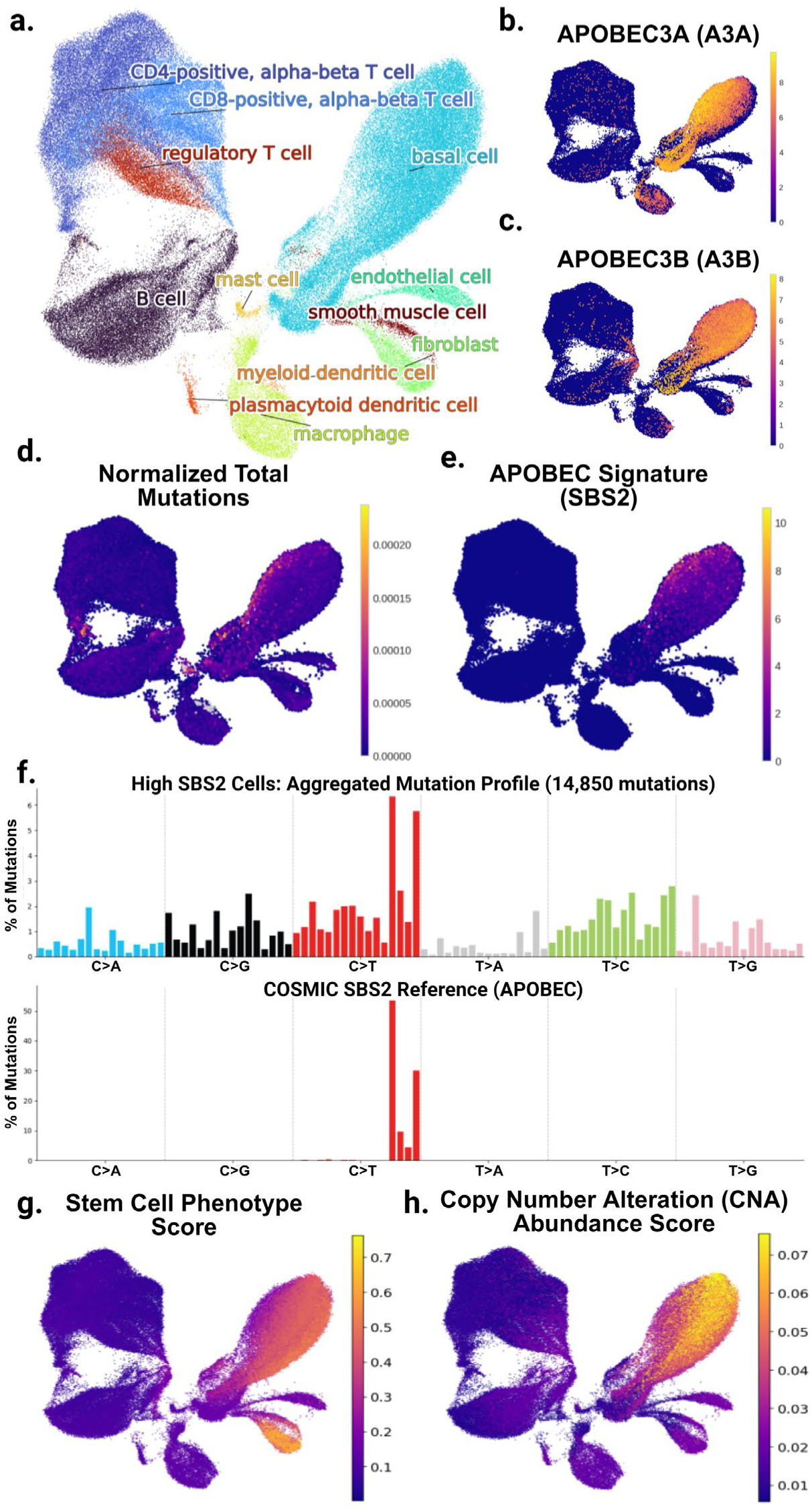
Single-cell resolution localizes SBS2 mutagenesis to basal epithelial cells co-expressing *A3A* and *A3B*. UMAPS of **(a)** cell type annotation of HNSCC single-cell data and **(b)** *A3A* and **(c)** *A3B* expression, respectively, showing enrichment in the basal cell cluster with partially distinct subregional localization. **(d)** Normalized somatic mutation count per cell (total SComatic-called mutations^25^ divided by callable sites with read depth ≥ 5), showing that basal epithelial cells carry the highest mutation burden. **(e)** SBS2 signature weight per cell from semi-supervised NMF refitting against COSMIC reference signatures^5^, co-localizing with the high-mutation basal cells. The region with the highest mutation burden and SBS2 weights co-localizes with the cells expressing elevated abundance of *A3A*, with variable to low expression of *A3B*. **(f)** Aggregated 96-trinucleotide-context mutation profile from the highest-SBS2 basal cells **(top)** compared to the pure COSMIC SBS2 reference signature **(bottom)**. The aggregated profile shows strong concordance with SBS2, dominated by C>T mutations in TCW contexts, with low-level background across other contexts consistent with single-cell data sparsity. UMAPS of **(g)** stem cell phenotype similarity score produced by CytoTRACE 2^26^ and **(h)** CNA abundance scores^27^ overlap and are enriched in basal cells with high expression of *A3B*.

**Figure 3.**
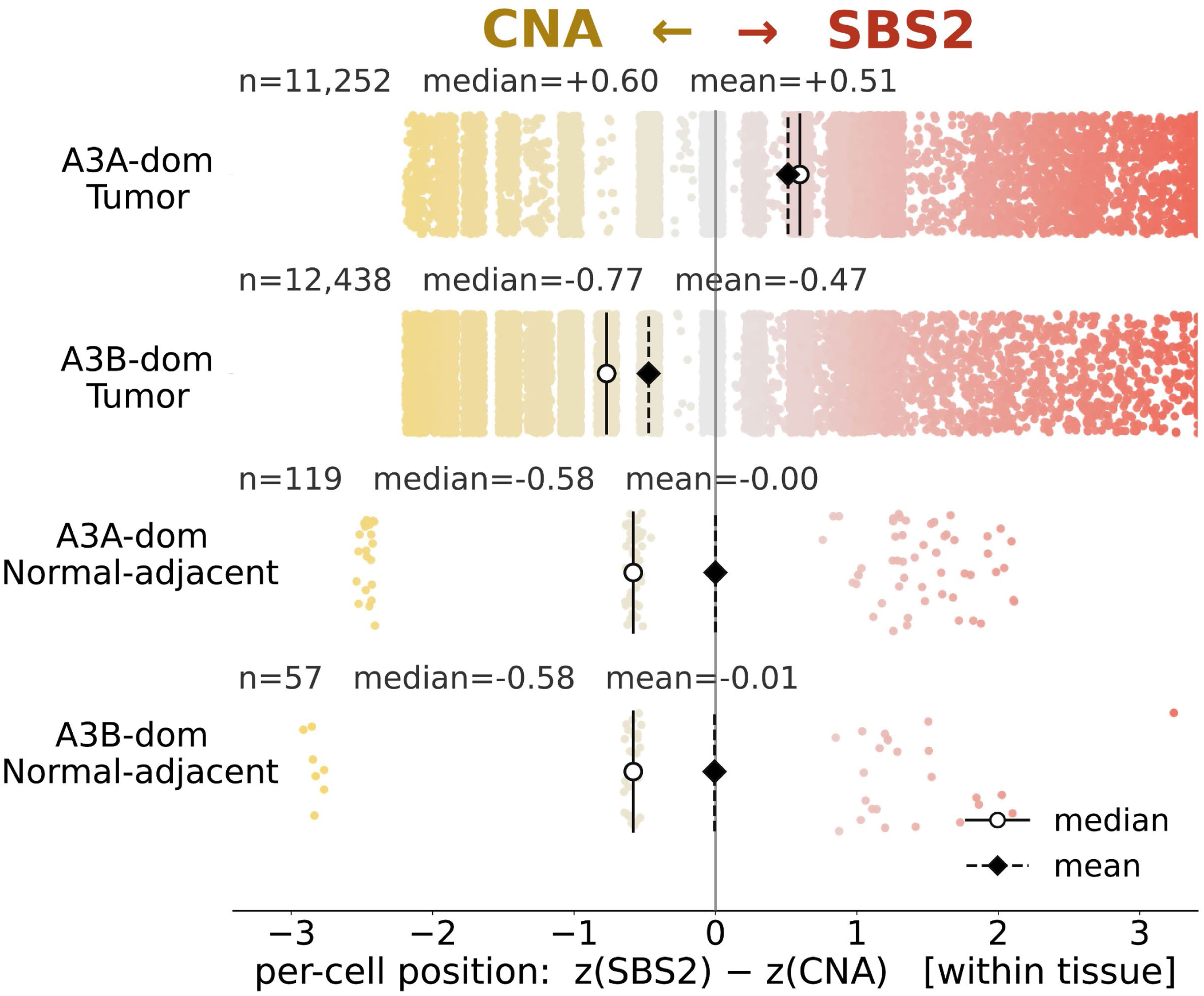
*A3A*– and *A3B*-dominant basal epithelial cells occupy opposite ends of an SBS2-versus-CNA axis in tumor but not in normal-adjacent tissue. Basal epithelial cells were classified by *A3* dominance, defined among cells expressing either enzyme as the fraction *A3A*/(*A3A* + *A3B*). Cells above 0.5 were termed *A3A*-dominant and below 0.5 *A3B*-dominant. Each point is one cell, placed along a within-tissue axis contrasting its standardized SBS2 burden against its standardized inferred CNA score. Cells to the right (red) are SBS2-enriched and low in CNA and cells to the left (yellow) are CNA-enriched. Rows show A3A-dominant and *A3B*-dominant basal epithelial cells separately in the tumor and in normal-adjacent tissue. Open circles mark the per-row median, filled diamonds the mean, and the vertical line the population center. In basal epithelial cells from tumor samples, *A3A*-dominant cells shift toward the SBS2 end and *A3B*-dominant cells toward the copy-number end (median +0.60, n = 11,252 and median –0.77, n = 12,438, respectively), whereas in normal-adjacent tissue the two groups overlap near a common center (n = 119 and n = 57). The separation is carried mainly by CNA. The *A3A* fraction correlates with the CNA score at Spearman rho = –0.44 in tumor versus +0.07 in normal-adjacent tissue, while its correlation with SBS2 is weak in both (+0.05 and +0.04). A small uniform horizontal jitter separated the discrete copy-number levels for display only; all medians, means, and correlations used unjittered values.

Among 31,912 basal epithelial cells with detected somatic mutations, 5,911 (18.5%) carried the A3-induced SBS2 mutational signature. The SBS2 signature burden in basal epithelial cells correlated more positively with the expression of *A3A* (Spearman rho = 0.149, p = 3.06 x 10^−158^) than *A3B* (rho = 0.052, p = 1.86 x 10^−20^). The stronger association with *A3A* is consistent with observations that *A3A* is the principal source of SBS2^6^, although both associations were relatively weak (Fig. 2d-e). The summed mutational profile of the cells with the highest SBS2 weights closely matched the COSMIC SBS2 reference, which is dominated by TCW>TTW mutations (Fig. 2f), supporting our deconvolution approach. Among SBS2-positive basal epithelial cells, 56.7% carried no detectable *A3A* and 49.8% carried no detectable *A3B*, with 32.1% lacking detectable expression of either enzyme. This might reflect episodic A3 expression^28^ or technical limits of single-cell RNA sequencing, including dropout of low-abundance transcripts. The second A3-associated signature, SBS13, was detected but carried the lowest mean weight of the fifteen COSMIC mutation signatures retained (mean weight 0.04, present in 7.7% of cells, against mean weight 0.19 and 18.5% for SBS2), was not enriched in basal epithelial cells, and showed no association with *A3A* expression (rho = 0.006, p = 0.32). We therefore focused on SBS2 as the readout of A3-driven mutagenesis.

We next asked whether basal epithelial cells with high A3-driven mutational burden also showed features of cancer progression, specifically transcriptional stemness and chromosomal instability. CytoTRACE 2 (RRID:SCR_029044)^26^ stemness scores and inferCNV (RRID:SCR_025804)^27^ CNA scores were both higher in basal epithelial cells relative to other cell types (Supp. Fig. 4). Neither feature co-localized with the highest SBS2 weights. Instead, a subpopulation of basal epithelial cells with high *A3B* expression showed elevated CNA and stemness (Fig. 2g-h). Across the compartment, *A3B* correlated positively with CNA (rho = 0.147, p = 9.85 x 10^−154^), whereas *A3A* correlated negatively (rho = –0.199, p = 3.35 x 10^−283^). Because *A3* induction is episodic^28^, expression level at capture is a noisy indicator of cumulative activity. We reasoned that the ratio between the two enzymes would be more stable. We therefore classified basal epithelial cells by *A3A* fraction, *A3A*/(*A3A* + *A3B*). Cells above 0.5 were *A3A*-dominant and cells below 0.5 were *A3B*-dominant. Each cell was then placed on a within-tissue axis contrasting standardized SBS2 burden with standardized CNA score. Scores were computed separately for tumor and normal-adjacent tissue (Fig. 3). In tumor tissue the two groups separated. *A3A*-dominant cells shifted toward the SBS2 end of the axis (median +0.60, n = 11,252) and *A3B*-dominant cells toward the CNA end (median –0.77, n = 12,438). In normal-adjacent tissue the two groups overlapped near a common center, although the small cell numbers limit interpretation (n = 119 and n = 57). CNA accounted for most of the separation. The *A3A* fraction correlated with the CNA score at rho = –0.44 in tumor versus +0.07 in normal-adjacent tissue, while its correlation with SBS2 was weak in both (+0.05 and +0.04). Together, these results point to two distinct associations between genomic alterations and *A3A*/*A3B* expression within the tumor epithelial compartment. *A3A* dominance is associated with SBS2, whereas *A3B* dominance is associated with CNA.

**Figure 4.**
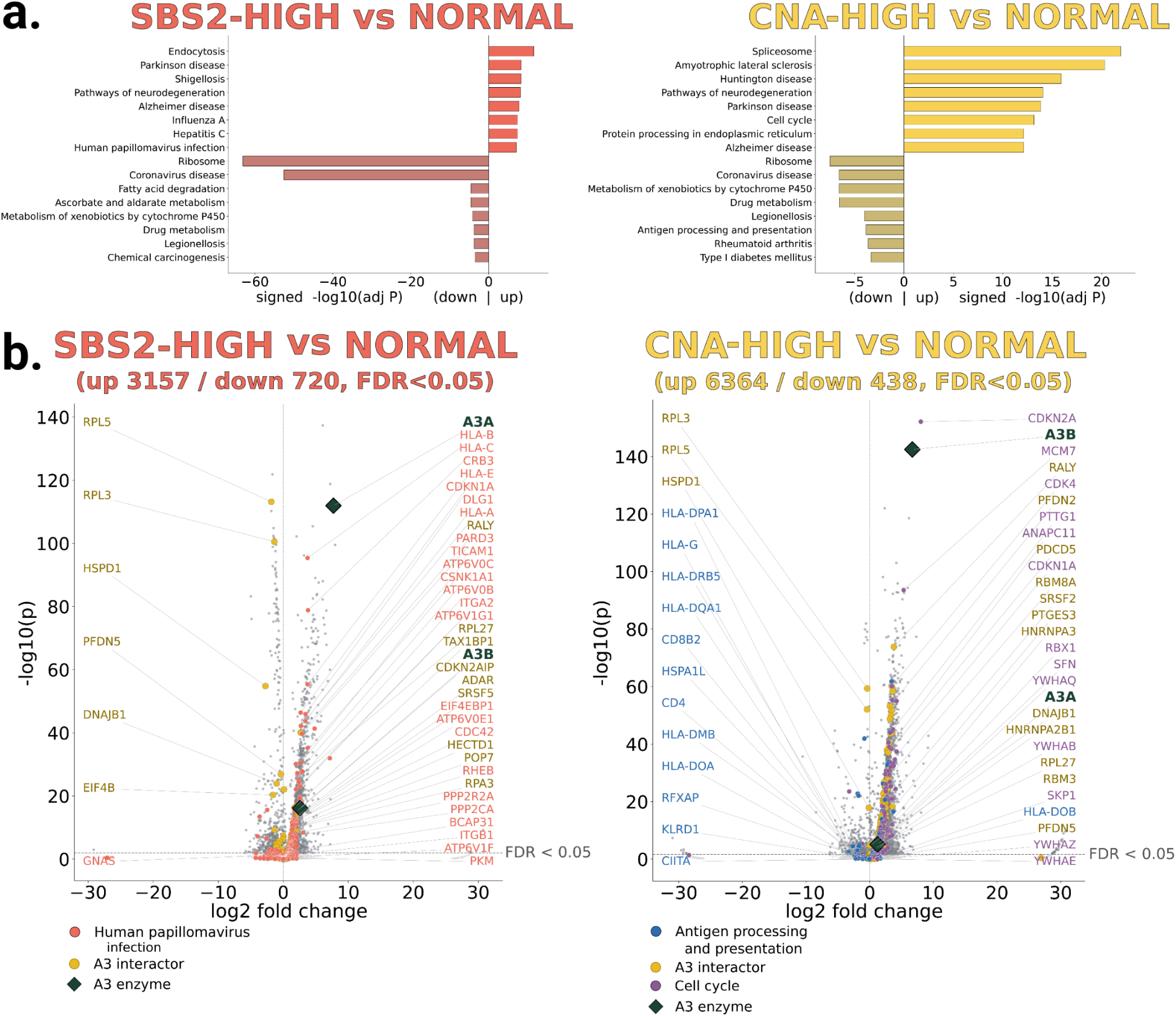
Differential expression separates the SBS2-HIGH and CNA-HIGH populations along an immune-recognition axis. **(a)** KEGG pathway enrichment on the differentially expressed genes of each tumor-versus-normal comparison, SBS2-HIGH versus NORMAL on the left and CNA-HIGH versus NORMAL on the right. Bars show signed –log10 adjusted p, with pathways enriched among up-regulated genes to the right of the zero line and pathways enriched among down-regulated genes to the left, eight per direction. **(b)** Volcano plots for the SBS2-/CNA-HIGH versus NORMAL comparison, plotting log2 fold change against –log10 P for every tested gene. Horizontal dashed line marks the FDR < 0.05 threshold. Genes belonging to the pathways of interest are colored: Human papillomavirus infection in the SBS2-HIGH panel, Cell cycle and Antigen processing and presentation in the CNA-HIGH panel. Catalogued A3 interactors are dark yellow and *A3A* and *A3B* are dark-green diamonds. Panel titles carry the number of up– and down-regulated genes at FDR < 0.05.

### 5.2 Single-cell network analysis identifies co-expression programs that may activate A3 mutagenic activity

To compare the two divergent tumor cell populations against cells from normal-adjacent tissue, we defined three equal-sized populations of 546 basal epithelial cells. SBS2-HIGH cells were drawn from tumor tissue and were enriched for SBS2 burden, expressed more *A3A* than *A3B*, and carried low CNA and stemness scores. CNA-HIGH cells were also drawn from tumor tissue and showed the reverse, with high CNA and stemness, more *A3B* than *A3A*, and no detectable SBS2. The NORMAL population was size-matched and drawn from normal-adjacent basal cells (see Methods). Cells were selected so that combined *A3A* + *A3B* expression was matched between the two tumor groups. Thus, the groups differed in which enzyme predominated rather than in their overall *A3* expression.

We first compared each tumor population with the NORMAL population by differential gene expression analysis (Fig. 4b). Both comparisons yielded extensive differences, with 3,877 genes differentially expressed in SBS2-HIGH cells and 6,802 in CNA-HIGH cells relative to NORMAL (false discovery rate, FDR < 0.05) (Supp. Files 1 and 2). SBS2-HIGH cells exhibited up-regulated viral-recognition and inflammatory programs, including the Kyoto Encyclopedia of Genes and Genomes (KEGG) (RRID:SCR_012773)^29^ pathways for human papillomavirus infection, influenza A, and hepatitis C (Fig. 4a). These pathways share substantial gene membership, so their co-enrichment represents one signal rather than three independent ones. By contrast, CNA-HIGH cells showed up-regulated cell cycle and down-regulated antigen processing and presentation genes (Fig. 4a). Thus, with respect to immune recognition, the CNA-HIGH cell population is more immunosuppressive, whereas the SBS2-HIGH population is more immune-visible. Despite these findings, differential expression alone does not show how these genes relate to *A3* enzymes, prompting us to examine differential co-expression networks.

We constructed differential co-expression networks comparing each of the SBS2-HIGH and CNA-HIGH populations with the NORMAL population, focusing on 174 previously catalogued A3 interactors that may function as A3 regulatory proteins^30,31^. Leiden community detection was used to identify clusters (modules) containing *A3A* and/or *A3B* genes^32^. In these differential networks, positive edges indicate co-expression that is stronger in tumor cells than in normal cells, and negative edges indicate the opposite. The SBS2-HIGH network (2,948 genes, 25 modules) and the CNA-HIGH network (4,886 genes, 38 modules) included 54 and 109 of the 174 interactors, respectively (Supp. Fig. 5). *A3A* and *A3B* fell within the same module in the SBS2-HIGH network but separated into distinct modules in the CNA-HIGH network. This suggests that the two enzymes belong to a shared co-expression program in cells with high SBS2, but to two separate programs in cells with high CNA.

**Figure 5.**
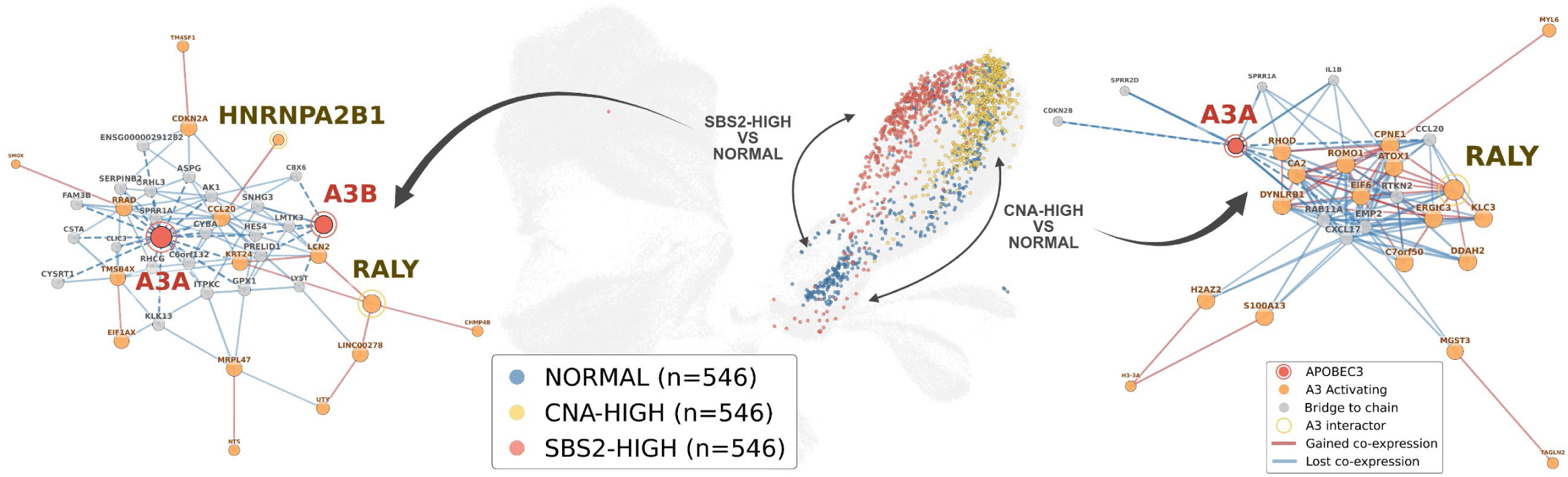
Single-cell differential co-expression network analysis identifies activating and inhibiting chains anchored by known A3 protein interactors. (Center) UMAP of 52,126 basal epithelial cells with three equal-sized populations (n = 546 each) highlighted: SBS2-HIGH (red), CNA-HIGH (yellow), and NORMAL (blue). Arrows indicate pairwise comparisons used to construct differential co-expression networks. (Left) Concordant chain subnetwork from SBS2-HIGH versus NORMAL. The six activating chains (orange nodes, red edges) contain the A3 interactors *RALY* and *HNRNPA2B1*, inflammatory (*CCL20, LCN2*), oxidative stress (*SMOX*), membrane remodeling (*CHMP4B*), and epithelial damage response genes (*KRT24, RRAD*). Dashed edges are the A3 wall: every edge linking chain genes to *A3A* or *A3B* (red nodes) is negative in the DIFF network, indicating co-expression stronger in normal tissue despite both enzymes being upregulated in the tumor. This is consistent with pulsatile A3 induction that degrades correlation structure in tumor relative to the stable, low expression in normal tissue. (Right) Concordant chain subnetwork from CNA-HIGH versus NORMAL. The 13-gene activating chain (orange nodes, red edges) also includes *RALY*, here within the A3A gene module, and is built from translation and metabolic genes (*CPNE1*, *EIFC*, *CA2*, *DYNLRB1*). *RALY* is present in the leading activating program in both tumor populations even though CNA-HIGH cells carry no SBS2 signature and *A3A* induction has fallen sharply (log2 fold change 1.2 versus 7.7 in SBS2-HIGH), marking it as a tumor-conserved activating program rather than one that tracks enzyme level. Dashed edges are the A3 wall: every edge linking A3B (red node) to its neighbors is negative in the DIFF network, co-expression stronger in normal tissue despite both enzymes being upregulated in tumor.

Both *A3A* and *A3B* were significantly up-regulated in tumor cells relative to normal cells in both network comparisons. Despite this induction, every edge connecting either enzyme to a first-degree neighbor was negative. Co-expression between the A3 enzymes and their immediate partners was therefore weaker in tumor cells than in normal cells throughout both networks. This formed a shell of uniformly anticorrelated edges surrounding each A3 enzyme, which we refer to as the “A3 wall”. This pattern is consistent with episodic A3 induction^28^, whereby expression rises sufficiently to generate SBS2 mutations but varies too greatly between cells to sustain the stable correlations observed in normal tissue. Because no gene was positively co-expressed with either enzyme in tumor cells, the sign of first-degree edges could not serve as a criterion for identifying candidate regulators.

Accordingly, we extended the analysis beyond the first-degree neighbors of *A3A* and *A3B* to the coherent transcriptional structure surrounding them. Chains of genes connected to one another by positive co-expression edges were traced outward from these *A3*-adjacent nodes and evaluated for concordant regulation. “Activating” chains were defined as those in which member genes were up-regulated in tumor cells and positively co-expressed with one another. “Inhibitory” chains were defined as those in which member genes were down-regulated and negatively co-expressed with one another in tumor cells. This approach identifies transcriptional programs surrounding the enzymes without requiring a positive edge directly to either enzyme.

Several activating chains contained catalogued A3 interactors. In the SBS2-HIGH network, *RALY* was a member of a six-gene activating chain comprising *LCN2*, *KRT24*, *LINC00278*, *CHMP4B* and *UTY*, and *HNRNPA2B1* was a member of a smaller activating chain that included *CCL20* (Fig. 5, left panel). In the CNA-HIGH network, *RALY* was a member of the largest chain within the *A3A* module, a 13-gene chain including *CPNE1*, *EIFC*, *CA2*, and *DYNLRB1* (Fig. 5, right panel). By contrast, inhibitory chains were present in both networks but contained no catalogued A3 interactor. The *A3B* module of the CNA-HIGH network consisted primarily of inhibitory chains. The largest inhibitory chain within the *A3A* module was a 70-gene squamous differentiation program comprising *MAB21L4*, *CYSRT1*, *SBSN*, and members of the *SPRR* family. *A3A* was positioned at the edge of this chain through connections to *SPRR1A*, *SPRR2D*, and *RAB11A*. Expression of these genes was higher in normal-adjacent tissue than in tumor cells, suggesting loss of the differentiation program that normally operates in the suprabasal compartment of squamous epithelium, consistent with the elevated stemness scores observed in CNA-HIGH cells.

Together, these analyses demonstrate that *A3* co-expression programs are extensively reorganized in tumor cells relative to normal tissue. Both enzymes are surrounded by uniformly negative edges, and the coherent programs adjacent to them separate into up-regulated activating chains and down-regulated inhibitory chains. Among the catalogued interactors, *RALY* was the only one present in an activating chain in both networks, making it the strongest candidate for a general activating role. The networks additionally contained genes showing stronger co-expression with *A3A* or *A3B* in tumor cells that have not previously been reported as A3 interactors (Supp. Files 3 and 4).

### 5.3 HPV16 life cycle stages are associated with distinct mutagenic programs in tumor epithelial cells

Two observations from the preceding analyses implicated HPV16 as the source of the divergence between the two tumor epithelial cell populations. First, in SBS2-HIGH cells viral-recognition programs are up-regulated relative to normal cells, whereas in CNA-HIGH cells the antigen processing and presentation pathway is down-regulated (Fig. 4). Second, innate immune and inflammatory genes including *LCN2* and *CCL20* appeared within the SBS2-HIGH co-expression network (Fig. 5). To assess the role of HPV16 directly, we examined the viral status of individual cells. Reads that did not map to the human genome were aligned to the HPV16 reference genome and quantified per cell. HPV16 reads were detected almost exclusively in our popV-annotated basal epithelial cells, which accounted for 94.6% of all HPV16-positive cells. An L-method breakpoint on raw HPV16 unique molecular identifier (UMI) counts defined the threshold above which viral read counts increased sharply. Cells above this threshold were classified as HPV16-positive, and lower counts were attributed to ambient contamination during library preparation, yielding 15,927 HPV16-positive basal epithelial cells (30.6%) (Fig. 6a-c).

**Figure 6.**
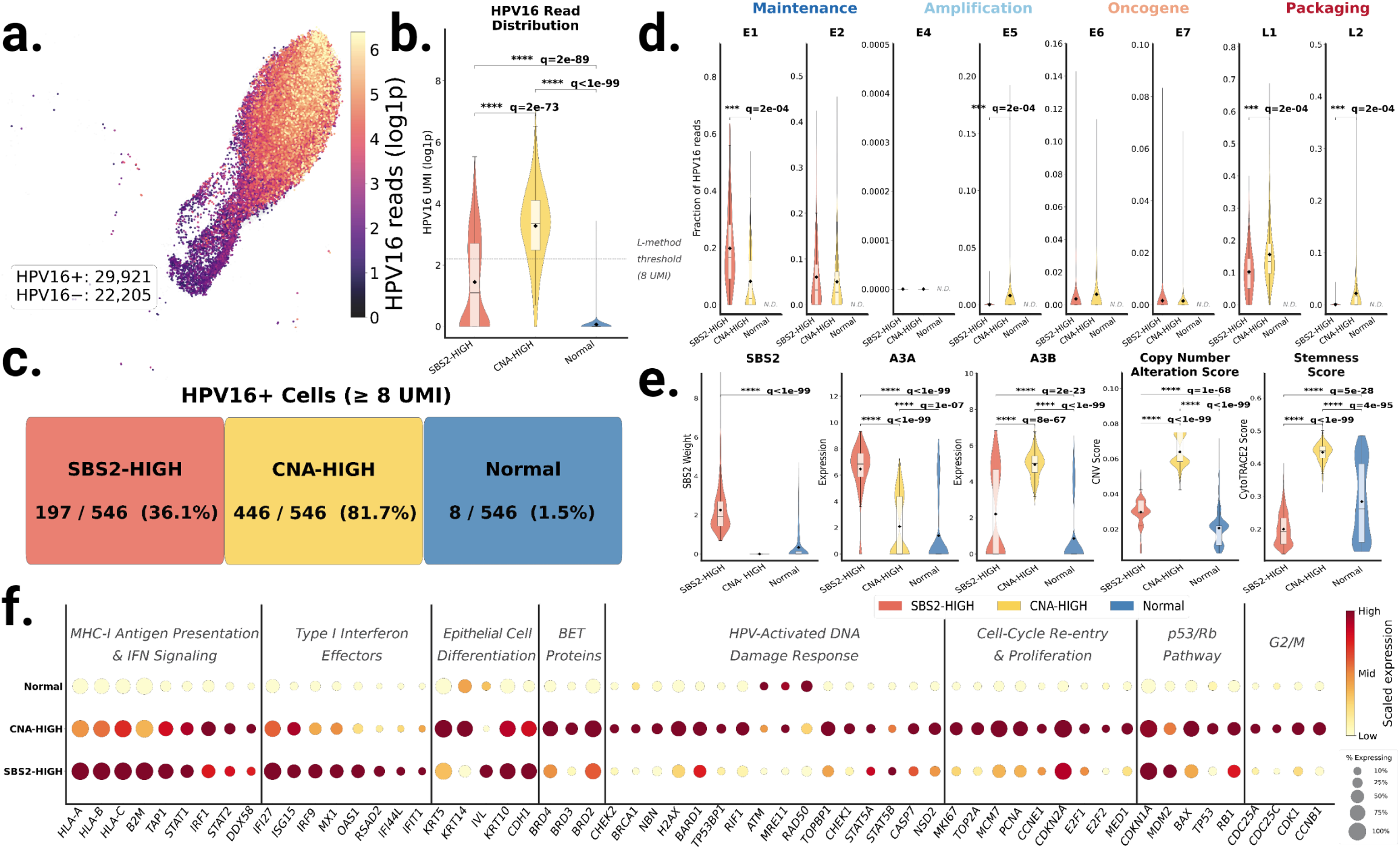
HPV16 life cycle stage is associated with the dominant A3 enzyme and the class of genomic damage in epithelial cells. **(a)** UMAP of epithelial cells colored by total HPV16 reads (log1p scale). HPV16 reads localize almost exclusively to this compartment (94.6% of all HPV16-positive cells). **(b)** Violin plot of HPV16 UMI read counts (log1p) per population. The dashed line indicates the L-method classification threshold (8 UMI). **(c)** Summary of HPV16-positive cell counts (≥ 8 UMI), per population. HPV16 positivity does not predict SBS2-HIGH membership (Fisher’s exact OR = 1.01, p = 0.91). **(d)** Mean per-cell fraction of total HPV16 reads mapping to each viral gene, grouped by viral gene function: maintenance (*E1*, *E2*), amplification (*E4*, *E5*), oncogene (*EC*, *E7*), and capsid (*L1*, *L2*). Restricted to HPV16-positive cells with alignment reads above zero. Brackets carry BH-adjusted q-values from a permutation test on the difference of means (10,000 permutations), corrected separately from the Mann-Whitney family in panels b and e. **(e)** Violin plots comparing SBS2 mutational signature weight, CNA score, transcriptional stemness, *A3A* expression, and *A3B* expression across SBS2-HIGH, CNA-HIGH, and NORMAL populations. Black diamonds indicate group means and white boxes show the interquartile range. Pairwise comparisons in panels b and e are Mann-Whitney U tests, BH-FDR corrected. **(f)** Host marker expression across 59 genes in eight functional tiers, ordered from the immune-visible maintenance end through differentiation to the viral assembly end: MHC-I antigen presentation and interferon signaling (9 genes), type I interferon effectors (8), epithelial cell differentiation (5), BET proteins (3), HPV-activated DNA damage response (16), cell-cycle re-entry and proliferation (9), p53/Rb pathway (5), and G2/M (4). Dot color is min-max scaled per gene across the three populations and dot size encodes the percentage of cells expressing the gene. The DNA damage response tier spans both arms, ATM (*ATM, CHEK2, NBN, MRE11, RAD50, BRCA1, H2AX, TP53BP1, RIF1, BARD1*) and ATR (*TOPBP1*, *CHEK1*), together with STAT5A and STAT5B, which act upstream of both arms in HPV-positive keratinocytes^35^. The tier also includes CASP7, the caspase that cleaves E1 downstream of CHK2^36^. NSD2, a double-strand break repair factor recruited by BRD4-L^37^, completes the tier. Total transcript levels of *ATM*, *MRE11*, and *RAD50* do not track activation of the HPV-associated DNA damage response, which is regulated by phosphorylation and complex assembly rather than by transcription.

Because progression of HPV-associated cancer tracks with the viral life cycle rather than with infection alone^33,34^, we asked whether life cycle stage explains the split between SBS2-HIGH and CNA-HIGH cells. We quantified per-cell viral reads across all HPV16 genes and grouped them into four functional categories^33,34^: maintenance (*E1*, *E2*), amplification (*E4*, *E5*), oncogene (*EC*, *E7*), and packaging (*L1*, *L2*). In all three populations, approximately two-thirds of HPV16 reads mapped to the non-coding upstream regulatory region (URR) rather than to a viral open reading frame (ORF) (63.3% in SBS2-HIGH, 67.1% in CNA-HIGH, and 64.6% in NORMAL cells). CNA-HIGH cells carried significantly more viral reads than SBS2-HIGH cells (Fig. 6b), averaging 2.6-fold more per HPV16-positive cell (235.1 versus 90.1, q = 1.7 x 10^−14^). As expected, only 8 of 546 NORMAL cells passed the HPV16-positive threshold, precluding statistical comparison. Life cycle contrasts were therefore restricted to SBS2-HIGH (n = 197) and CNA-HIGH (n = 446) cells.

Among HPV16-positive cells, viral gene expression placed the two tumor populations at different stages of the HPV16 life cycle (Fig. 6d). Maintenance-stage genes accounted for a larger share of viral reads in SBS2-HIGH than in CNA-HIGH cells (25.9% versus 13.2%, maintenance category q = 1.3 x 10^−4^), a difference driven by *E1* (q = 2.0 x 10^−4^). Virus assembly showed the reverse, with capsid reads higher in CNA-HIGH than in SBS2-HIGH cells (17.8% versus 10.3%, packaging category q = 1.3 x 10^−4^; *L1* and *L2*, gene q = 2.0 x 10^−4^ for each). CNA-HIGH cells also carried a greater share of *E5* reads (0.80% versus 0.01%, q = 2.0 x 10^−4^), consistent with active viral genome amplification. Oncogene output was minor in both states, with less than 1% of viral reads mapping to *EC* and *E7* (0.53% in SBS2-HIGH and 0.79% in CNA-HIGH). *E2* did not differ between the populations (q = 0.11), and we found no evidence of *E2* loss that accompanies viral integration, consistent with both populations carrying episomal virus.

Host gene expression aligned with these viral stages (Fig. 6e,f). We profiled 59 host genes across eight functional tiers, of which 47 differed significantly between SBS2-HIGH and CNA-HIGH cells. SBS2-HIGH cells, enriched for the viral maintenance stage, were immune-visible and differentiating. Major histocompatibility complex (MHC) class I antigen presentation was elevated (*B2M* q = 9.0 x 10^−100^, *HLA-A* q = 7.5 x 10^−42^, *HLA-B* q = 1.2 x 10^−15^, *HLA-C* q = 5.9 x 10^−8^, *TAP1* q = 1.0 x 10^−4^). All eight type I interferon effectors peaked in these cells, seven of them significantly (*IFI27* q = 2.9 x 10^−48^, *OAS1* q = 2.3 x 10^−30^, *RSAD2* q = 3.0 x 10^−28^, *IRFS* q = 1.2 x 10^−15^, *MX1* q = 1.3 x 10^−13^, *IFI44L* q = 4.4 x 10^−7^, *IFIT1* q = 7.8 x 10^−3^; *ISG15* q = 0.20). By contrast, the upstream sensing and signaling components of the same axis did not separate the two tumor populations (*STAT1*, *IRF1*, *STAT2*, and *DDX58*). This indicates that these populations differ in interferon effector output rather than in pathway engagement. Involucrin (IVL), a terminal differentiation marker, was likewise higher in SBS2-HIGH cells than in either other population (2.68 versus 0.09, q = 5.2 x 10^−70^). As expected from the selection criteria, *A3A* expression exceeded *A3B* in these cells, and the magnitude of that difference was substantial (6.46 versus 2.21) (Fig. 6e). Several genes from the SBS2-HIGH activating chains belong to this same inflammatory program. *CCL20* is co-expressed with *HNRNPA2B1*, *LCN2* is a neighbor of *RALY*, and *SMOX* occupies a separate chain with *RRAD* (Fig. 5, left panel). The presence of these genes alongside elevated interferon effector output suggests that *A3A* induction occurs within an active inflammatory context. However, because expression was measured at a single timepoint, these data cannot order the induction of *A3A* relative to the inflammatory program.

CNA-HIGH cells enriched for the viral assembly stage showed a different program. The DNA damage response required for HPV16 genome amplification was elevated (*CHEK2* q = 1.4 x 10^−5^, *BRCA1* q = 1.1 x 10^−8^, *NBN* q = 1.3 x 10^−3^, *H2AX* q = 4.3 x 10^−11^). The BRD4-recruited repair factors *TP53BP1* (q = 2.4 x 10^−6^) and *RIF1* (q = 6.6 x 10^−11^) were elevated alongside them, as was the ATR arm of the response, represented by *TOPBP1* (q = 8.4 x 10^−14^) and *CHEK1* (q = 7.2 x 10^−11^). The bromodomain and extra-terminal motif (BET) proteins were also elevated in CNA-HIGH cells (*BRD2* 4.29 versus 5.00 q = 1.6 x 10^−9^, *BRD3* 0.64 versus 1.90 q = 1.8 x 10^−25^, and *BRD4* 2.56 versus 3.48 q = 1.3 x 10^−11^). The increase was largest for BRD3, which is the BET family member required for HPV16 genome amplification, whereas BRD2 is dispensable^37^. Phosphorylated BRD4 recruits TP53BP1 and BARD1 to the HPV16 origin of replication during differentiation-associated genome amplification, in a BRD4 isoform-specific manner, with recruitment shifting from TP53BP1 toward BARD1 as amplification proceeds^37^. That recruitment is governed by phosphorylation rather than transcript abundance, and consistent with this, *BARD1* did not differ between the two populations (q = 0.076). The DNA damage response was also accompanied by G2/M machinery (*CDC25A* q = 2.4 x 10^−16^, *CDC25C* q = 3.0 x 10^−12^, *CDK1* q = 4.7 x 10^−18^, *CCNB1* q = 9.8 x 10^−27^). The active DNA damage response provides the repair factors that HPV16 uses to amplify its genome. *CASP7* was also elevated (q = 5.7 x 10^−3^), consistent with the caspase cleavage of E1 required for genome amplification^36^. Cell-cycle re-entry accompanied these changes, with 8 of the 9 genes in that tier significantly up-regulated in CNA-HIGH cells (*MCM7* q = 1.7 x 10^−47^, *TOP2A* q = 1.2 x 10^−25^, *MKIC7* q = 2.4 x 10^−19^, *PCNA* q = 3.8 x 10^−13^, *MED1* q = 1.8 x 10^−8^, *E2F1* q = 1.1 x 10^−5^, *E2F2* q = 5.1 x 10^−5^, *CCNE1* q = 1.4 x 10^−4^). As expected from the selection criteria, *A3B* expression exceeded *A3A* in these cells (4.95 versus 2.08) (Fig. 6e), and they retained a proliferative basal identity (*KRT14* q = 8.9 x 10^−57^, *KRT5* q = 4.4 x 10^−7^). In line with a previous report^38^, cell-cycle phase followed the same division. SBS2-HIGH cells were predominantly in G1 (63.9%), whereas CNA-HIGH cells were predominantly in S or G2/M (81.7%, with 41.8% in G2/M) (Supp. Fig. 6a). The same separation was observed when all basal cells were grouped by *A3* enzyme dominance or by *A3* presence and absence rather than by population (Supp. Fig. 6b).

These findings led us to reconsider the relationship between cell state and viral stage in the two populations. In epithelial cells from normal-adjacent tissue, the dominant A3 enzyme matched the host cell state described previously^18,38^ for squamous epithelium, with *A3A* in differentiating cells and *A3B* in cycling cells (Supp. Fig. 7). The viral stage did not follow the same pattern. Genome maintenance canonically occurs in the basal and parabasal compartment and virus assembly in differentiating suprabasal cells^34^, yet SBS2-HIGH cells carried maintenance-stage virus while expressing *IVL*, and CNA-HIGH cells carried assembly-stage virus while retaining a proliferative basal keratin profile (elevated *KRT5* and *KRT14*) with *IVL* nearly absent. Cluster-level annotation by popV assigned both populations to the basal compartment and did not distinguish these two epithelial states. This is consistent with *E6* and *E7* uncoupling the transcriptional profile from the position along the differentiation trajectory. SBS2-HIGH and CNA-HIGH cells accordingly appear to represent two epithelial states along a maintenance-to-assembly axis, distinguished by viral gene usage rather than by differentiation status. Together, these results support the association of *A3A* expression with the genome maintenance stage and *A3B* expression with the virus assembly stage of the HPV16 life cycle. The HPV16 life cycle also appears to influence the type of genomic damage that accumulates, with SBS2 representing A3-associated mutational burden and CNA representing chromosomal instability. The SBS2:CNA ratio may therefore serve as a marker of a tumor cell’s position along the maintenance-to-assembly axis of the HPV life cycle, providing a molecular readout with potential implications for treatment stratification.

**Figure 7.**
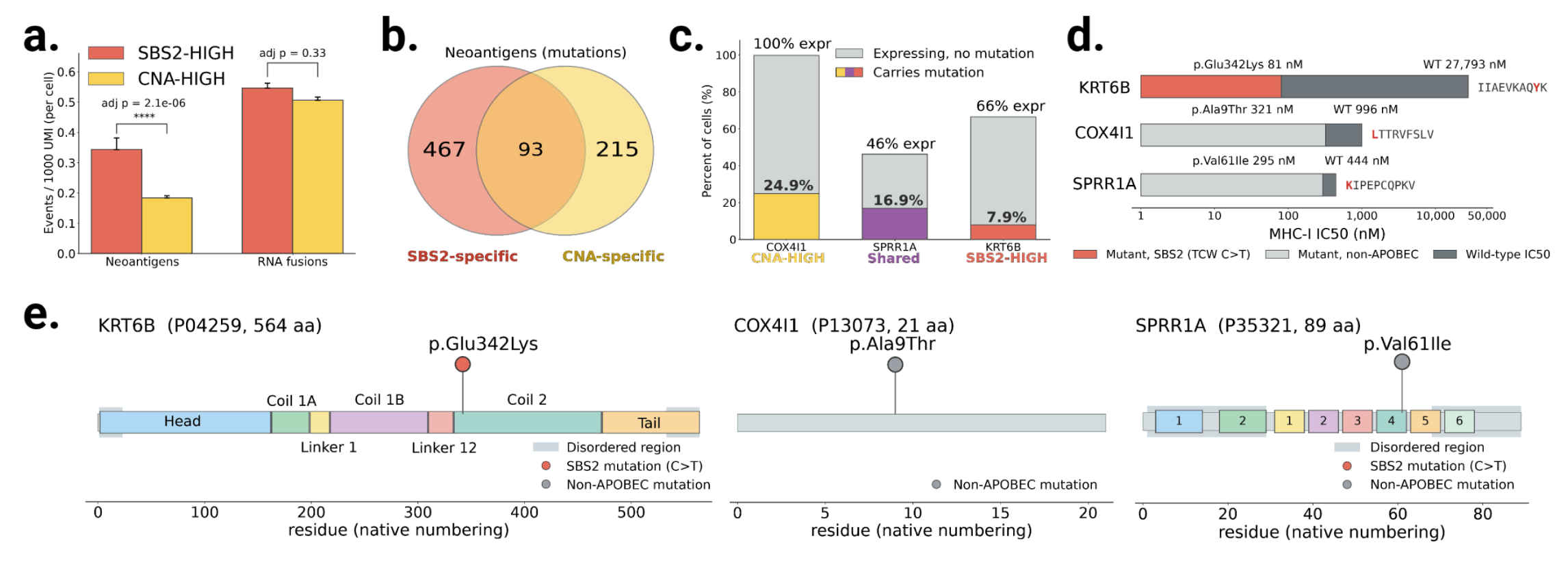
Neoantigen landscape of the immune-visible and immune-evasive epithelial cell populations and prevalence-prioritized vaccine candidates. (a) Neoantigen-forming mutation burden and RNA-fusion burden per cell, normalized per 1000 UMI to control for sequencing depth, in SBS2-HIGH versus CNA-HIGH cells. (b) Venn diagram of the overlap of neoantigen-forming mutations between the two populations. (c) Expression and mutation prevalence for the three prioritized candidates, COX4I1 (CNA-specific), SPRR1A (shared), and KRT6B (SBS2-specific). Bar height is the percentage of cells expressing the gene, and the colored base is the percentage of cells carrying the neoantigen-forming mutation (its clonal prevalence), on a population-consistent denominator (specific candidates over their own 546 cells, the shared candidate over the pooled 1,092). (d) MHC-I binding of the wild-type and mutant peptide for each featured neoantigen, shown as the wild-type baseline binding and the binding gain conferred by the mutation (lower indicates stronger binding). Predicted IC50 values are annotated: COX4I1 Ala9Thr, 996 to 321 nM; SPRR1A Val61Ile, 444 to 295 nM; KRT6B Glu342Lys, 27,793 to 80.5 nM. The mutated residue is shown in red within each peptide sequence. Bars are colored by mutational context (clean T[C>T]W SBS2/APOBEC). (e) Position of each featured neoantigen-forming mutation on its protein, with functional domains and regions annotated from UniProt in the native transcript numbering. Lollipops mark the mutated residue, colored by mutational context.

### 5.4 Patient-specific contributions to each mutagenic fate track with viral load, viral life cycle direction, and the fraction of cells expressing the matching *A3* enzyme.

Donor-specific effects are a well-established source of variation in single-cell datasets^39^. To determine whether individual patients contributed disproportionately to either the SBS2-HIGH or CNA-HIGH population, we stratified both populations by donor and compared each patient’s contribution against the size of their epithelial cell pool. The 546 SBS2-HIGH cells originated from 11 of 14 patients, and three contributed more cells than their epithelial compartment share predicted (chi-square p = 4.46 x 10^−309^). Patients SC013 (221 cells, 40.5%, 6.9-fold), SC029 (133 cells, 24.4%, 2.1-fold), and SC001 (50 cells, 9.2%, 2.5-fold) together accounted for 404 of the 546 cells in the SBS2-HIGH population (74.0%) (Supp. Fig. 8a). The CNA-HIGH population was similarly concentrated in two patients (chi-square p < 1 x 10^−300^). Patients SC027 (285 cells, 52.2%, 8.9-fold) and SC001 (163 cells, 29.9%, 8.2-fold) together accounted for 82.1% of that population (Supp. Fig. 8a). Only SC001 contributed to both. The two fates therefore arise predominantly in different tumors, indicating that the divergence is determined at least partly at the patient level rather than solely at the individual-cell level.

To assess whether the network results depended on these donors, we reconstructed the differential networks after excluding one patient at a time. Because the activating and inhibitory chains differ between the SBS2-HIGH and CNA-HIGH networks, no common chain set exists across both, and this analysis was therefore applied to the SBS2-HIGH network and its contributing donors. Removal of any single high contributor reduced network size but left the A3 wall intact, with activating-chain genes and module structure largely preserved (Supp. Fig. 9). The co-expression program is therefore shared across donors rather than specific to any one of them. The networks were therefore similar across patients, whereas the capacity of individual tumors to generate either state differed, implicating viral and microenvironmental factors.

Although the two tumor populations differed in overall HPV16 abundance, we found that viral load at the patient level did not predict which fate a tumor contributed to. Library-normalized HPV16 load associated with contribution to both fates at nearly the same strength (Spearman rho = +0.506 for SBS2-HIGH and +0.513 for CNA-HIGH), and two patients carrying high load, SC010 and SC022, contributed to neither population. Viral burden nonetheless appeared to establish a minimum threshold. Every high contributor carried HPV16 at more than 0.1% of its epithelial-compartment transcripts, though three further patients (SC010, SC022, and SC026) also exceeded that threshold without contributing disproportionately to either population. Viral burden therefore appeared necessary for a tumor to contribute cells to either population but did not determine which fate predominated. Fate was predicted by the distribution of viral reads rather than their abundance. The life cycle enrichment observed within the SBS2-HIGH and CNA-HIGH populations held at the patient level, with every high contributor carrying more than 40% of its viral coding reads in the stage matching its fate: maintenance genes (*E1* and *E2*) for the SBS2-HIGH contributors, and late-stage genes (*E4*, *E5*, *L1*, and *L2*) for the CNA-HIGH contributors (Supp. Fig. 8b).

We next asked whether patient-level differences in *A3A* and *A3B* expression accounted for these contributions. The high contributors were not the highest expressors of either enzyme, and SC001 ranked mid-cohort for *A3A* (mean 1.12, rank 6 of 14) (Supp. Fig. 8c). SC005 carried the highest *A3A* expression in the cohort at 3.03 yet contributed to SBS2-HIGH at 0.7-fold. This is consistent with our earlier observation that *A3A*/(*A3A* + *A3B*) dominance, rather than raw *A3* expression, was more strongly associated with the accumulation of SBS2 or CNA (Fig. 3). Across the cohort, per-cell *A3* output was near-constant, varying 1.4-fold for *A3A* and 1.2-fold for *A3B*, whereas the fraction of cells expressing them varied 76.5-fold and 342.3-fold, respectively. Prevalence rather than level therefore distinguished the high contributors in an enzyme-specific manner. *A3A* prevalence tracked SBS2-HIGH contribution (rho = +0.581, p = 0.030) but not significantly with CNA-HIGH contribution (rho = +0.142, p = 0.63), whereas *A3B* prevalence tracked with CNA-HIGH contribution (rho = +0.660, p = 0.010) but not significantly with SBS2-HIGH contribution (rho = +0.400, p = 0.16).

Taken together, viral burden, viral life cycle direction, and broad expression of the matching *A3* enzyme all appear necessary to shift a tumor toward one fate. Individual tumors illustrated why none is sufficient alone (Supp. Fig. 10). SC013 and SC027 carried identical viral loads, 0.17% of epithelial-compartment transcripts each, yet contributed to opposite fates, with SC013 enriched for maintenance-stage virus and *A3A* expression and SC027 enriched for late-stage virus and *A3B* expression. SC005 carried the highest *A3A* expression in the cohort but only a third as much virus and contributed to neither population. SC010 carried abundant maintenance-stage virus but expressed *A3A* at the cohort median and likewise contributed to neither. SC001 carried the most virus of any tumor and contributed to both. Applying all three criteria jointly separated every high contributor from the remainder of the cohort (Fisher exact p = 0.0027 for SBS2-HIGH and p = 0.011 for CNA-HIGH). A tumor therefore contributed more cells than expected to a mutational fate when three conditions were met: HPV16 accounted for at least 0.1% of its epithelial-compartment transcripts, at least 40% of its viral coding reads fell within the matching life cycle stage, and the matching *A3* enzyme was expressed in at least one in five epithelial cells for *A3A* or one in two for *A3B*. These thresholds were derived from a cohort of 14 patients and will require validation in larger cohorts.

### 5.5 Neoantigen landscape and therapeutic target identification

The two epithelial states differ not only in the mutations they carry but in whether those mutations can be presented to the immune system. Tumors accumulate somatic mutations that can produce neoantigens^40,41^. Because they are tumor-specific, neoantigens are attractive targets for cancer vaccines, including multivalent mRNA vaccines that prime T-cell responses against several epitopes simultaneously^42^. The efficacy of such a strategy depends on the immune state of the tumor. Immune-visible tumors maintain active antigen presentation and T-cell infiltration and tend to respond to immunotherapy, whereas immune-evasive tumors down-regulate antigen presentation and largely escape T-cell surveillance^43^. SBS2-HIGH cells expressed antigen-presentation genes at higher levels than CNA-HIGH cells, marking the two populations as immune-visible and immune-evasive, respectively. Beyond immune recognition of HPV16 itself, we reasoned that heavily mutated cells might also carry a greater burden of neoantigens, which would contribute independently to an immune-visible state. To assess this, we called somatic protein-altering variants in each population. We then predicted MHC-I binding for the mutant and wild-type peptides across ten common HLA class I alleles and compared the resulting neoantigen and RNA-fusion burden on a per-cell basis (Fig. 7a).

SBS2-HIGH cells produced more predicted neoantigen-forming mutations than CNA-HIGH cells (560 versus 308, 1.82-fold, p = 9.50 x 10^−18^). Because SBS2-HIGH cells were sequenced less deeply, we also compared burden normalized to 1,000 UMI, and the excess remained (0.343 versus 0.184 neoantigens per 1,000 UMI, Benjamini-Hochberg-adjusted p = 2.15 x 10^−6^; Fig. 7a). RNA fusions, by contrast, did not follow the same division. Fusion burden was statistically indistinguishable between the two populations both per cell and per UMI (0.547 versus 0.506 fusion junctions per 1,000 UMI, Benjamini-Hochberg-adjusted p = 0.327; Fig. 7a). Raw junction counts were marginally higher in CNA-HIGH than in SBS2-HIGH cells (5,566 versus 5,128) and highest in normal-adjacent basal cells (6,625, 12.1 per cell), indicating that fusion burden does not distinguish the two immune states.

We next examined how these neoantigen-forming mutations partitioned between the two populations (Fig. 7b). Across both populations we identified 775 unique neoantigen-forming mutations, of which 93 were shared, 467 were SBS2-HIGH-specific, and 215 were CNA-HIGH-specific. SBS2-HIGH-specific neoantigens arise in cells with an active antigen-presentation program and may therefore be more likely to elicit immune responses, although their utility may be restricted to immune-visible tumors. CNA-HIGH-specific neoantigens, by contrast, arise in immune-evasive cells and are less likely to drive a functional T-cell response, but they remain of interest because targets for immune-evasive tumor cells are otherwise scarce.

Shared neoantigen mutations are the most broadly applicable, as they are present regardless of the HPV16 life cycle stage of the tumor cell. These mutations occurred in PI3, SERPINB2, and KRT6A, with SPRR1A Val61Ile representing the most clonally prevalent shared candidate (Fig. 7c,d). A subset of the shared mutations (15 of 93) showed patterns consistent with immune selection in CNA-HIGH cells. In each case, the gene was either disrupted by a CNA-HIGH RNA fusion that removed the mutant junction, or transcriptionally silenced such that the mutant protein was not produced. Both patterns are also consistent with the genomic instability and transcriptional dysregulation that characterize CNA-HIGH cells, and the present data cannot distinguish immune selection from these alternatives. Neoantigens arising at HLA-A, HLA-B, and HLA-C were excluded from this count, because these hyperpolymorphic loci are prone to alignment artifacts and the apparent binding gains occurred at prevalences approaching that of the population as a whole.

To prioritize candidates, we ranked neoantigen-forming mutations by the fraction of tumor cells carrying them (Supp. File 5), which identified three leading candidates, one from each partition (Fig. 7c,d). COX4I1 Ala9Thr is a CNA-HIGH-specific neoantigen carried in 24.9% of CNA-HIGH cells and expressed nearly universally (99.8%). SPRR1A Val61Ile is a shared neoantigen carried in 16.9% of tumor cells. KRT6B Glu342Lys is an SBS2-HIGH-specific neoantigen carried in 7.9% of SBS2-HIGH cells. Of the three, KRT6B Glu342Lys showed the largest predicted MHC-I binding gain, converting a non-binding wild-type peptide (predicted half-maximal inhibitory concentration IC50 27,793 nM) into a strong binder (81 nM), whereas COX4I1 Ala9Thr and SPRR1A Val61Ile are more prevalent but show lower predicted binding strength.

KRT6B Glu342Lys is of particular interest because it arises in a clean T[C>T]W context characteristic of SBS2, linking a high-affinity neoantigen to the A3 mutational process characteristic of the immune-visible population. Together, these three candidates indicate how each partition of the mutational landscape might be addressed: SPRR1A Val61Ile as a shared target spanning both immune states, KRT6B Glu342Lys for the immune-visible SBS2-HIGH population, and COX4I1 Ala9Thr for the immune-evasive CNA-HIGH population. Because prioritization was based on the fraction of cells carrying each mutation rather than on expression level alone, each candidate is positioned to engage a substantial share of the population it targets.

## 6 Discussion

Expression of both *A3A* and *A3B* is elevated in HPV16-positive HNSCC, yet expression level alone accounts for little of the variation in A3-driven mutational burden. Bulk tissue cannot resolve this question because it averages across both non-tumor and tumor cells, as well as tumor cells at different stages of the viral life cycle. By quantifying gene expression and mutational signatures in individual single cells from HPV16-positive tumors and matched normal-adjacent tissue, we found that HPV16 life cycle plays a key role in A3-induced mutagenesis. As an infected cell progresses from maintenance toward virus assembly, the dominant A3 enzyme and its regulatory network shift, changing the resulting pattern of genomic damage.

The breakdown of normal A3 regulation in tumor cells is evident in the co-expression networks. Comparison of each tumor population against normal-adjacent tissue revealed A3 interaction patterns that bulk tissue obscures. Our differential network analysis placed *A3A* and *A3B* among up to 109 of the 174 catalogued A3 interactors^30,31^ within densely connected gene modules. Within this network, we traced an activating chain containing the interactors *RALY* and *HNRNPA2B1*, which suggests that catalogued A3 binding partners may act as regulatory proteins that modulate mutagenic activity rather than as passive associates. The same chains contained secreted inflammatory mediators. *CCL20* was co-expressed with *HNRNPA2B1*, and *LCN2* with *RALY*, placing *A3A* induction within an active inflammatory program.

The clearest indication of altered regulation was the A3 wall. Although both *A3A* and *A3B* were up-regulated in tumor cells, every edge connecting either enzyme to its neighbors was stronger in normal tissue than in tumor tissue. In normal epithelium, the enzymes are expressed at low levels in tight coordination with their partners, and in tumor cells that coordination is lost. Considered alongside the SBS2-positive cells that carried no detectable A3 expression at capture, the pattern is consistent with pulsatile A3 induction, in which transient bursts leave permanent mutations without generating stable correlations. Episodic A3 mutagenesis has been described in cell-line models^28^, and our data extend that observation to primary tumors at single-cell resolution.

The stage of the viral life cycle determines both the amount and the type of damage. SBS2-HIGH cells carried virus in maintenance, with *E1*-enriched viral expression, immune-visible antigen presentation, elevated interferon effector expression, and *A3A* dominance. CNA-HIGH cells carried virus in the assembly stage, with capsid expression, a DNA damage response spanning both the ATM and ATR arms, G2/M machinery, and *A3B* dominance. A cell’s position within the life cycle therefore appears to determine which *A3* enzyme predominates and which class of damage accumulates: SBS2 under *A3A* during the HPV16 maintenance stage and chromosomal instability under *A3B* during the virus assembly stage.

In normal-adjacent epithelial tissue, we observed that *A3B* expression is associated with proliferating epithelial cells and *A3A* with differentiating epithelial cells, in agreement with previous work^18^. This pattern is altered by HPV16 in subpopulations of the tumor. SBS2-HIGH cells remained consistent with normal-adjacent tissue, since a differentiating, *IVL*-expressing, *A3A*-dominant cell is where *A3A* is expected. CNA-HIGH cells are the departure. The viral assembly program normally unfolds in differentiating and terminally differentiated epithelial cells, yet here it operated in cells that retained a proliferative basal keratin identity (*KRT5* and *KRT14* elevated) with *IVL* nearly absent, a decoupling consistent with *E6* and *E7* sustaining cell cycle entry while suppressing differentiation. Because popV reference-based annotation assigns cells by transcriptional profile, both populations were labeled basal epithelial despite occupying different positions along this axis. We therefore regard SBS2-HIGH and CNA-HIGH not as distinct cell types but as two epithelial states along a maintenance-to-assembly axis. The SBS2:CNA ratio may accordingly indicate a tumor’s position along this axis, providing a potential molecular measure of life cycle progression. This interpretation assumes that the rate at which SBS2 is converted to CNA is broadly comparable across tumors, which remains to be established.

The divergence between these epithelial states also operates at the level of the patient. Three patients supplied 74.0% of the SBS2-HIGH cells and two supplied 82.1% of the CNA-HIGH cells, with only one tumor contributing to both, indicating that the two fates arise largely in different tumors. What distinguished these patients was not the level of *A3A* or *A3B* expression, since the highest expressor in the cohort contributed to neither fate. Mean expression among expressing cells varied at most 1.4-fold across patients, whereas the fraction of cells expressing the enzyme at all varied by more than an order of magnitude, and it was this prevalence that tracked fate. Patient-level *A3* dysregulation is therefore a penetrance phenomenon, determined by the fraction of cells expressing *A3* rather than by the magnitude of expression in individual cells. Together, these findings define a set of tumor-level prerequisites involving viral load, life cycle direction, and *A3* expression prevalence. A cell reaches either mutational state only where the patient environment supplies virus, the matching life cycle stage, and an enzyme expressed across much of the compartment.

These prerequisites also shape what the immune system can detect. Maintenance-stage cells carried 1.82-fold more predicted neoantigen-forming mutations than viral assembly-stage cells, an excess that persisted after normalization for sequencing depth, while retaining antigen presentation. The higher neoantigen burden therefore occurs in the population with greater antigen-presentation capacity. Of the 775 unique mutations, 93 were shared between the two states, and 15 of those showed patterns consistent with removal in CNA-HIGH cells through fusion disruption or transcriptional silencing. Ranking by clonal prevalence rather than expression identified one candidate per partition: SPRR1A Val61Ile as a shared target, KRT6B Glu342Lys for the immune-visible population, and COX4I1 Ala9Thr for the immune-evasive population. KRT6B Glu342Lys arises in a clean T[C>T]W context characteristic of SBS2 and converts a non-binding wild-type peptide into a strong binder, linking a prevalent target to the mutational process characteristic of the immune-visible state.

Several limitations temper these conclusions. The central limitation of this work is the number of tumors underlying the patient-level findings. Additionally, single-timepoint capture means that our evidence for pulsatile A3 induction is inferential rather than direct. Differential co-expression identifies association rather than causation, so the chains described here indicate candidate A3 activators that require functional testing. Two of these results are in apparent tension. Leave-one-patient-out reconstruction left the A3 wall and the activating chains intact, whereas the contribution analysis shows that the SBS2-HIGH and CNA-HIGH populations derive largely from a small number of patients. The two are not in conflict, because they measure different quantities. The reconstruction measures the resilience of the co-expression program in cells that already reached the SBS2-HIGH state, and that program persisted when any one of the three high contributors was removed. By contrast, the contribution analysis measures how many cells from each patient reach that state, which varied markedly between tumors. The co-expression program therefore appears to be shared across patients. We propose that the number of cells entering either state depends instead on a permissive patient environment defined by viral load, life cycle stage, and the fraction of epithelial cells expressing the matching A3 enzyme. However, this hypothesis rests on three tumors for SBS2-HIGH and two for CNA-HIGH. Testing this model will require a substantially larger cohort. A central question is whether HPV16 consistently establishes a microenvironment that promotes both HNSCC development and subsequent immune evasion. Finally, MHC-I binding for every neoantigen reported here was algorithmically predicted rather than measured. Both the binding itself and any resulting T-cell response require experimental validation before these candidates can be regarded as vaccine targets.

## 7 Methods

### 7.1 Single-cell acquisition, processing, and mutational analysis

Single cell reads were aligned to GRCh38 with Cell Ranger (RRID:SCR_017344)^44^, and cells were filtered in adaptive mode by median-absolute-deviation (MAD) outlier detection. This was applied to both the log-transformed total and gene counts (more than 5 MADs from the median), with additional removal of cells above 20% mitochondrial reads, genes detected in fewer than 20 cells, and Scrublet-flagged doublets (RRID:SCR_018098)^45^. Cell types were annotated with popV^23^ using the Tabula Sapiens reference (RRID:SCR_022314)^46^ on raw counts with batch correction on sample identity. Final calls were made at the Leiden-cluster level by summing normalized popV confidence per cluster, which suppresses individual misclassifications. Somatic mutations were called with SComatic^25^, which pools reads within each annotated cell type, requires a variant in at least 5 cells with adequate coverage, filters against a panel of normals and known RNA-editing sites, removes any variant seen in more than one cell type as likely germline, and normalizes each cell’s mutation count to its callable sites. The 96-trinucleotide-context matrix was decomposed against HNSCC-relevant COSMIC^5^ v3.4 signatures by semi-supervised non-negative matrix factorization (NMF). The deconvolution always retains SBS2, SBS13, and SBS5 and adds other COSMIC signatures by scree-plot elbow on reconstruction error, with per-cell weights stored in the AnnData object. All of these analysis steps were performed as part of our custom analysis pipeline called ClusterCatcher (RRID:SCR_029058), released with this publication.

### 7.2 SBS2 and CNA association analysis with *A3A* and *A3B* expression

To relate *A3* expression to mutational fate (Fig. 3), epithelial cells were partitioned into tumor and normal-adjacent tissue by sample source. Per-cell SBS2 was taken from the NMF weights (mapped by cell barcode), *A3A* and *A3B* from the log-normalized expression matrix, and per-cell CNA burden from the inferCNV score (inferCNVpy^27^, window 250, CytoTRACE 2-defined normal cells as reference). *A3* dominance was defined for each cell expressing either enzyme (*A3A* + *A3B* > 0) as the fraction *A3A*/(*A3A* + *A3B*), with values above 0.5 classified *A3A*-dominant and below 0.5 *A3B*-dominant. For each cell expressing *A3A* or *A3B* (*A3A* + *A3B* > 0), we placed it on a single axis contrasting its point-mutation burden against its copy-number burden. SBS2 weight and CNA score were each rank-transformed and standardized (z-scored) across this expressing population, separately within tumor and within normal-adjacent tissue, and the cell’s position was its standardized SBS2 rank minus its standardized CNA rank. Standardizing within each tissue places SBS2 and CNA on a common scale before subtraction and prevents overall magnitude differences between tumor and normal-adjacent cells from driving the axis. Positive values therefore indicate SBS2-leaning cells and negative values CNA-leaning cells. We then related *A3* dominance to each burden by Spearman correlation of the *A3A* fraction with SBS2 weight and with CNA score, computed separately in tumor and normal-adjacent cells.

### 7.3 Differential expression, pathway enrichment, and differential co-expression network analysis

Differential expression for each tumor-versus-normal comparison used scanpy’s (RRID:SCR_018139)^47^ rank_genes_groups function with a Wilcoxon rank-sum test on the log-normalized matrix. Genes detected in fewer than 10 cells were excluded. P-values were Benjamini-Hochberg corrected and genes at adjusted p < 0.05 were retained. Up– and down-regulated sets were then tested separately for KEGG^29^ pathway enrichment (KEGG_2021_Human) through the Enrichr (RRID:SCR_001575)^48^ API via GSEApy (RRID:SCR_025803)^49^, at an adjusted p < 0.05 threshold. The background for each test was the set of genes tested in that comparison rather than the whole genome.

For the network, three non-overlapping basal epithelial populations of 546 cells each were defined. The group size of 546 was set from an inflection point in the ordered SBS2 weight distribution across tumor epithelial cells, above which SBS2 weight rose sharply. This count fell close to the 554 basal epithelial cells available from normal-adjacent samples, permitting size-matched selection. To extend this initial finding while incorporating the observation that SBS2 burden was more strongly associated with *A3A* expression dominance and CNA with *A3B* expression dominance (Fig. 3), we built a composite scoring system to select the 546 cells in each tumor population that best represented the distinct states along the *A3A:*SBS2 and *A3B*:CNA axes, with a size-matched selection from normal-adjacent basal cells. SBS2-HIGH cells were selected from SBS2 > 0 cancer-tissue cells by a composite score weighting SBS2 weight (40%), inverse CNA score (20%), inverse CytoTRACE 2 stemness (20%), and *A3A*/(*A3A*+*A3B*) expression fraction (20%). CNA-HIGH cells were selected from SBS2 = 0 cancer-tissue cells by a composite weighting CNA score (40%), total *A3* (*A3A*+*A3B*) expression range-matched to the SBS2-HIGH mean by Gaussian proximity (20%), CytoTRACE 2 stemness (20%), and *A3B*/(*A3A*+*A3B*) expression fraction (20%). Within each candidate pool, each selection metric was converted to a percentile rank (0 to 1) and inverted where low values were favored (i.e. low CNA, low stemness in SBS2-HIGH), the resulting terms were combined as a weighted sum and the top 546 cells were retained. The NORMAL population was a random sample of 546 cells from normal-adjacent tissue (546 of the 554, 98.6% available basal cells).

Spearman correlation matrices were computed independently in each population and subtracted to form a differential matrix (DIFF = HIGH minus reference). The DIFF threshold was selected dynamically per comparison by a maximum fragmentation-rate criterion, among thresholds retaining at least one connection for both A3A and A3B, breaking ties toward the lower and more inclusive value. Thresholds were lower than is typical for bulk co-expression, because single-cell correlation magnitudes are attenuated. Community structure was identified with the Leiden algorithm on the full disconnected graph, with edge weights set to |delta-rho|. Resolution was chosen by a sweep scored on modularity x ARI x evenness, followed by a component-aware merge that preserves satellite modules corresponding to whole small components. Recovery of known A3 interactors was assessed against 174 interactors compiled from published affinity-purification datasets^30,31^.

Activating and inhibiting chains were defined by directional coherence, in which a gene’s differential expression direction agrees with the sign of its DIFF edges. Positive DIFF edges indicate co-expression gained in the tumor condition and negative edges co-expression maintained in the reference. Two coherent subgraphs were built per comparison: an activator subgraph of positive edges among up-regulated nodes, and an inhibitor subgraph of negative edges among down-regulated nodes, with *A3A* and *A3B* included in both as reachability targets. Because *A3A* and *A3B* fell into different modules in the CNA-HIGH versus NORMAL network, each enzyme’s module was analyzed separately. Within each *A3*-containing module, connected components of the coherent subgraphs were enumerated as candidate chains and annotated for membership of known A3 interactors.

### 7.4 HPV16 life cycle analysis

Per-cell HPV16 abundance was taken from Kraken2^50^ raw UMI counts (recovered from the sparse feature-barcode matrices), and an L-method breakpoint at 8 UMIs split cells into negative (0 to 7), and positive (UMIs ≥ 8). To resolve life cycle stage and specific viral genes expressed, reads that did not map to the human genome were aligned to the HPV16 reference (NC_001526.4) with splice-aware minimap2^51^. Genes were grouped into four functional groups following established life cycle biology^33,34^: maintenance (*E1*, *E2*), amplification (*E4*, *E5*), oncogene (*EC*, *E7*), and packaging (*L1*, *L2*); the amplification signal reflects *E5*, because the spliced *E1^E4* transcript does not map to its standalone coordinates. Because HPV16-positive CNA-HIGH cells carried roughly 2.6-fold more viral reads than SBS2-HIGH cells, per-gene counts were expressed as fractions of total HPV16 reads to avoid biasing the life cycle comparison by viral load, retaining the URR (about two-thirds of reads). Per-gene and per-group fractions were compared by a permutation test on the difference of means, Benjamini-Hochberg corrected within each family, while population-defining and viral-load distributions used Mann-Whitney with figure-wide correction.

### 7.5 Epithelial differentiation state scoring

Epithelial differentiation states were scored across the basal epithelial compartment using the four states and marker genes defined for healthy tonsil epithelium^18^. Per-cell scores were computed with scanpy score_genes against 50 control genes per expression bin. In normal-adjacent cells, scores were compared between *A3A*-dominant (n = 119) and *A3B*-dominant (n = 57) cells by two-sided Mann-Whitney U tests, Benjamini-Hochberg corrected across the four states, and each cell was assigned to the state with the highest score after standardizing the four scores within that population. No assignment was made in tumor cells, because the states were defined in uninfected epithelium.

### 7.6 Patient contribution and network robustness

To test whether the SBS2-HIGH population was a conserved program or driven by individual patients, we compared each patient’s observed SBS2-/CNA-HIGH cell count to the count expected if SBS2-/CNA-HIGH membership were proportional to that patient’s share of the epithelial compartment. A chi-squared test of independence on a patient by group-membership contingency table was used for this comparison. Patients contributing more than twofold the number of cells expected from their epithelial cell share were designated high-contributing. Network generalizability was tested by leave-one-patient-out re-runs across the three SBS2-HIGH high-contributing patients, scoring preservation of activating-chain genes, within-community gene agreement (adjusted Rand index), overall gene overlap (Jaccard), and *A3*-wall integrity (percentage of connections between *A3A* or *A3B* and neighboring nodes that are stronger in normal tissue).

Three per-patient determinants were then related to contribution. Viral burden was computed as summed HPV16 UMIs over summed total UMIs across a patient’s epithelial cells, reported as a percentage of the epithelial-compartment transcriptome. Life cycle stage balance was computed on HPV16-positive cells only, as summed maintenance or viral late-stage reads over summed coding reads, pooled across the patient’s cells with the URR excluded. Patients with fewer than 10 HPV16-positive cells were not assigned a stage. *A3A* and *A3B* were summarized two ways: as the mean log-normalized expression across all tumor epithelial cells, and as the percentage of those cells with nonzero expression. A per-patient mean taken across all cells assigns zero to non-expressing cells, so it decomposes into the fraction of expressing cells multiplied by the mean among expressing cells. Both terms were computed separately, allowing a threshold on the mean to be expressed equivalently as a fraction of expressing cells.

### 7.7 Neoantigen and fusion analysis

SComatic variants were formatted into per-population (SBS2-HIGH, CNA-HIGH, and NORMAL) VCFs, annotated with SnpEff (RRID:SCR_005191)^52^ (GRCh38.p14). SBS2-/CNA-HIGH VCFs were background subtracted against NORMAL to remove neoantigens that were observed in the normal epithelial cell population from tumor results. Because SComatic calls variants within the epithelial-cell compartment, this leaves a somatic, cell-type-restricted set. Trinucleotide context for each variant was read from the GRCh38 reference genome, and mutations in a TCW context (where W = A or T) were classified as APOBEC-associated, with C>T changes corresponding to SBS2 and C>G to SBS13. For each missense variant, mutated and wild-type peptides of 8 to 11 residues were built on the Ensembl release 115 proteome (RRID:SCR_002344)^53^ through a multi-layer transcript, gene-symbol, alias, gene-ID, and position-offset lookup that verified the wild-type residue at each step (98.6 to 98.7% mapping). Peptides were scored with MHCflurry (RRID:SCR_028577)^54^ across 10 common HLA class I alleles (A*01:01, A*02:01, A*03:01, A*24:02, B*07:02, B*08:01, B*35:01, B*44:02, C*04:01, C*07:01). Population coverage for this allele set was estimated with the Immune Epitope Database (IEDB) Population Coverage tool (RRID:SCR_006604)^55^, giving 90.5% coverage of the world population, the fraction of individuals carrying at least one allele in the panel.

Each neoantigen was then localized in single cells. Using the single-cell genotype table, we counted the fraction of tumor cells carrying the neoantigen-forming mutation (its clonal prevalence) on an SBS2-/CNA-HIGH unique or shared denominator. Population-specific neoantigens were scored over their own 546 cells and shared neoantigens over the pooled 1,092 cells, and candidates were ranked by clonal prevalence together with the MHC-I binding gain (wild-type minus mutant IC50). With the neoantigen-forming mutations that were shared between both groups, we flagged neoantigens showing patterns consistent with immune escape in CNA-HIGH cells through either expression silencing (>2-fold downregulation) or fusion disruption of the gene. HLA-A/B/C neoantigens were retained in the binding analysis but excluded from this escape count because their hyperpolymorphic loci are prone to mapping artifacts and their apparent binding gains occurred at prevalences approaching that of the population as a whole. Neoantigen and fusion burden were compared between the two populations per cell, normalized per 1000 UMI to control for the lower depth of SBS2-HIGH cells, and corrected together by the Benjamini-Hochberg procedure. For fusions, FASTQs were re-aligned with STARsolo (RRID:SCR_021542) in chimeric mode, and chimeric junctions passed a confidence cascade (unique mapping, alignment-score and gene-boundary filters, protein-coding partners, and artifact-locus exclusion). Junctions detected in the NORMAL population were subtracted from both tumor populations, removing germline and recurrent non-somatic junctions. Recurrent group-exclusive pairs were identified, and each population’s neoantigen genes were tested against the other’s exclusive fusion partners to gauge directional immune escape.

### 7.8 Data availability

Single-cell data from HPV16-positive HNSCC tumors and matched normal samples were obtained from the gene expression omnibus (GEO) database with accession number (GSE173468). All analyses were performed in Python 3.11. All analysis code along with Anaconda virtual machine environments used to run the analysis are available at https://github.com/Diako-Lab/Network-Architecture-of-APOBEC3-Driven-Mutagenesis-in-HPV-Head-and-Neck-Cancers. Raw reads were retrieved from the Sequence Read Archive via GEO accession with SRAscraper (https://github.com/Diako-Lab/SRAscraper)(RRID:SCR_029060) and processed through the ClusterCatcher single-cell pipeline (https://github.com/Diako-Lab/ClusterCatcher), both available in the Diako-Lab GitHub repository.

## 8 Acknowledgements

We thank Dr. Gregory C. Ippolito (Texas Biomedical Research Institute) and Dr. Sophia Liu (Broad Institute of MIT and Harvard and Ragon Institute of Mass General Brigham, MIT, and Harvard) for reviewing the text and providing comments. J.D.L. received support from the Forum Foundation Fellowship (Texas Biomedical Research Institute: 1519801). C.-M.C. is presently supported by US Department of War Breast Cancer Research Program Breakthrough Award HT9425-26-1-E445, NIH grant 1R01CA288743-01A1, and the Chung-Ho Chen Cancer Research Fund. D.E. is presently supported by the NIH grants 5R01AI179465-03 and 5R01AI189230-02. This work received computational support from UT San Antonio’s High Performance Computing Center (HPCC), Arc, operated by Tech Solutions. This work also received computational support from the Texas Biomedical Research Institute HPC. Figures were created with BioRender.com.

## 9 Declaration of generative AI and AI-assisted technologies in the manuscript preparation process

During the preparation of this work, the authors used Claude and ChatGPT for proofreading and for generating code used in data analysis and figure production. The author(s) reviewed and edited the output as needed and take full responsibility for the content of the published article.

---

**Supplemental Figure 1:**
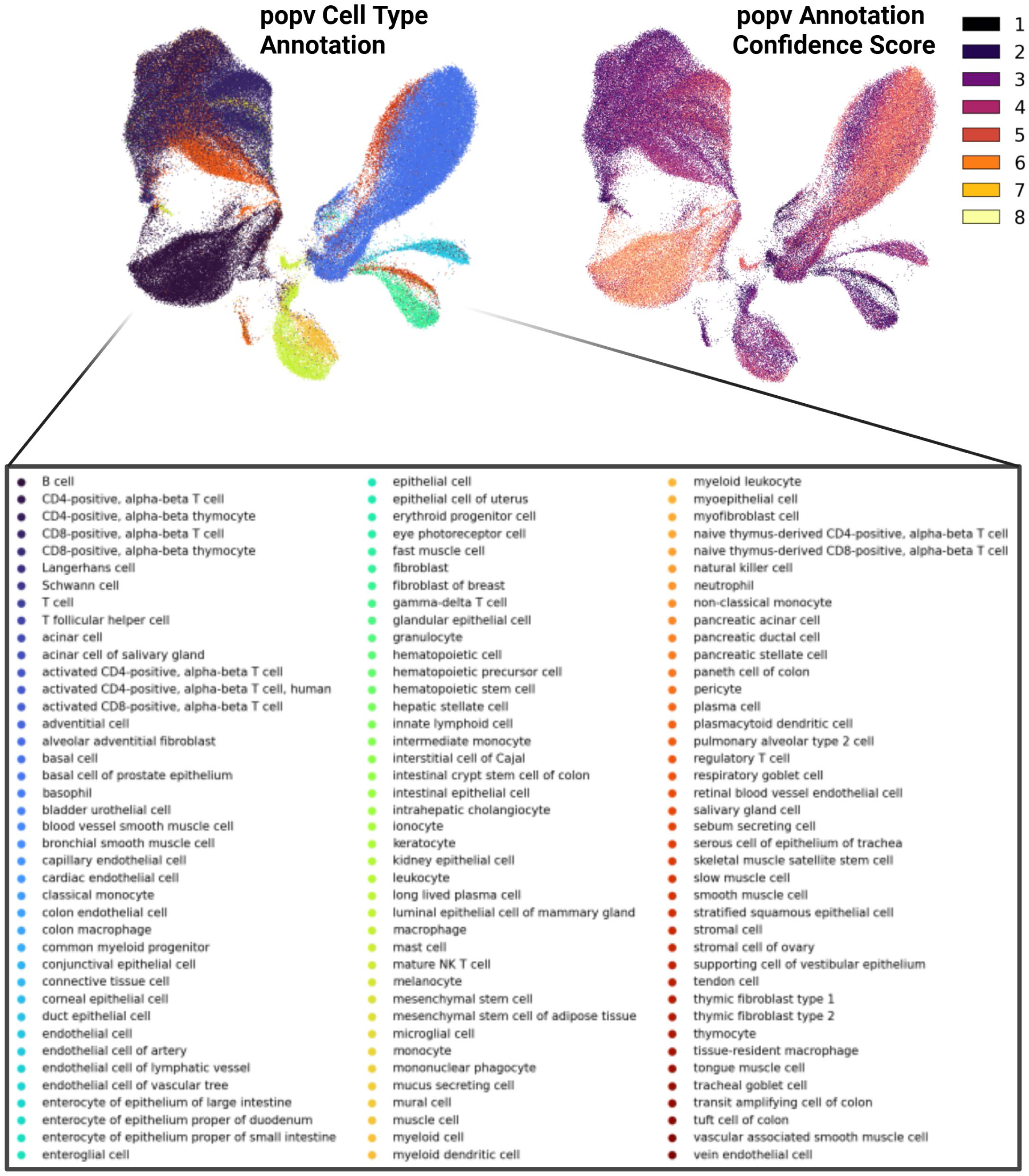
Per-cell popV cell type predictions in HNSCC single-cell data. UMAP embedding of all cells colored by individual popV cell type predictions^23^ using the Tabula Sapiens reference model^46^. Each cell is assigned the cell type with the highest popV prediction probability, independent of cluster membership. While the majority of cells cluster by predicted cell type, residual misclassification is visible at cluster boundaries, particularly among low-confidence predictions. These scattered misassigned cells motivate the cluster-level annotation refinement used in Fig. 1, 2a, in which each Leiden cluster is assigned a single cell type identity based on the highest summed normalized popV confidence score across all cells within the cluster, effectively filtering out low-confidence individual assignments while preserving the dominant cell type signal.

**Supplemental Figure 2.**
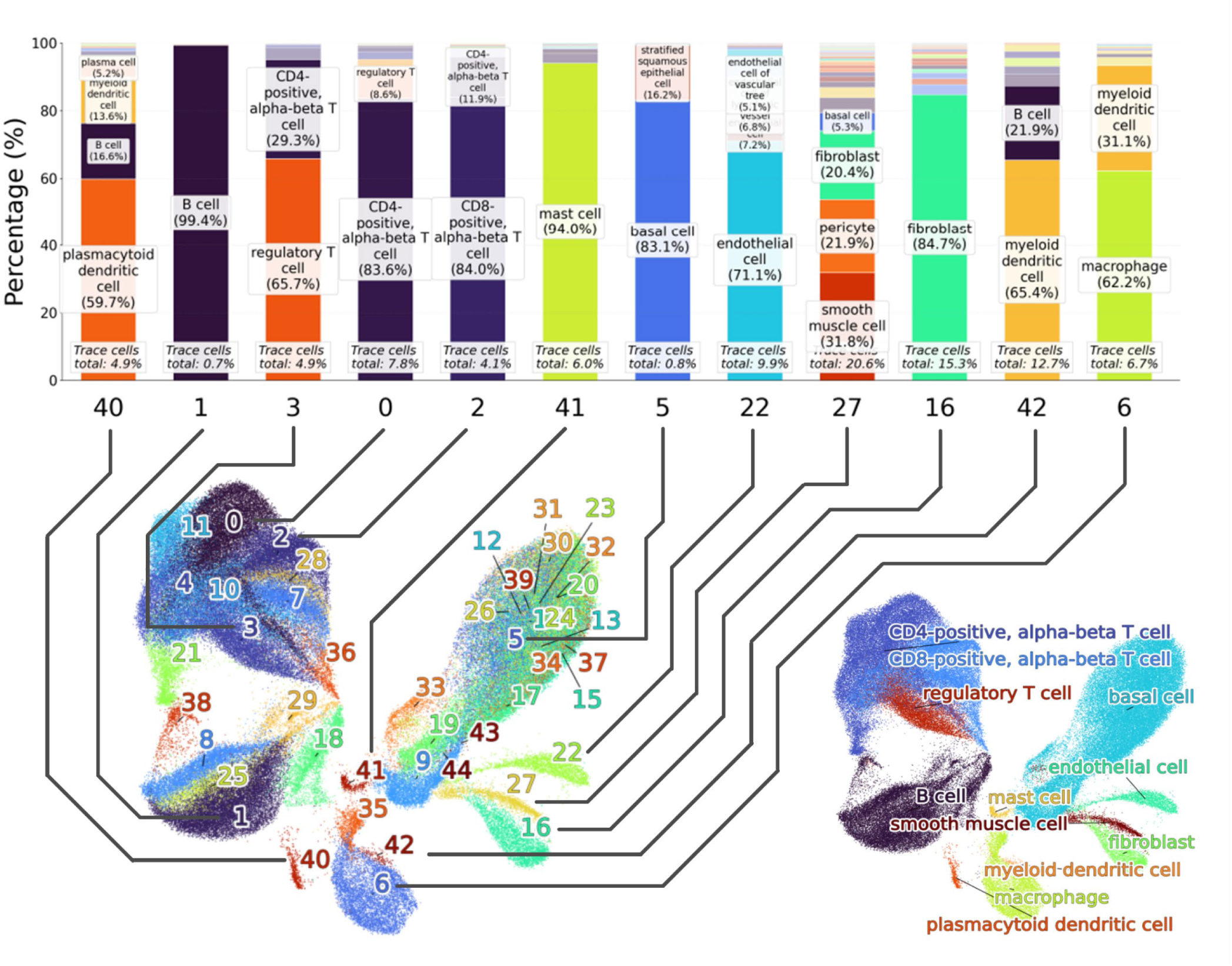
Cell type annotation of HNSCC single-cell data. **Left:** stacked bar plot showing the proportion of popV-predicted cell types within each Leiden cluster, demonstrating that clusters are dominated by a single cell type. **Right:** UMAP with final cell type assignments determined by the highest summed normalized popV confidence score per cluster, resolving 12 cell populations.

**Supplemental Figure 3.**
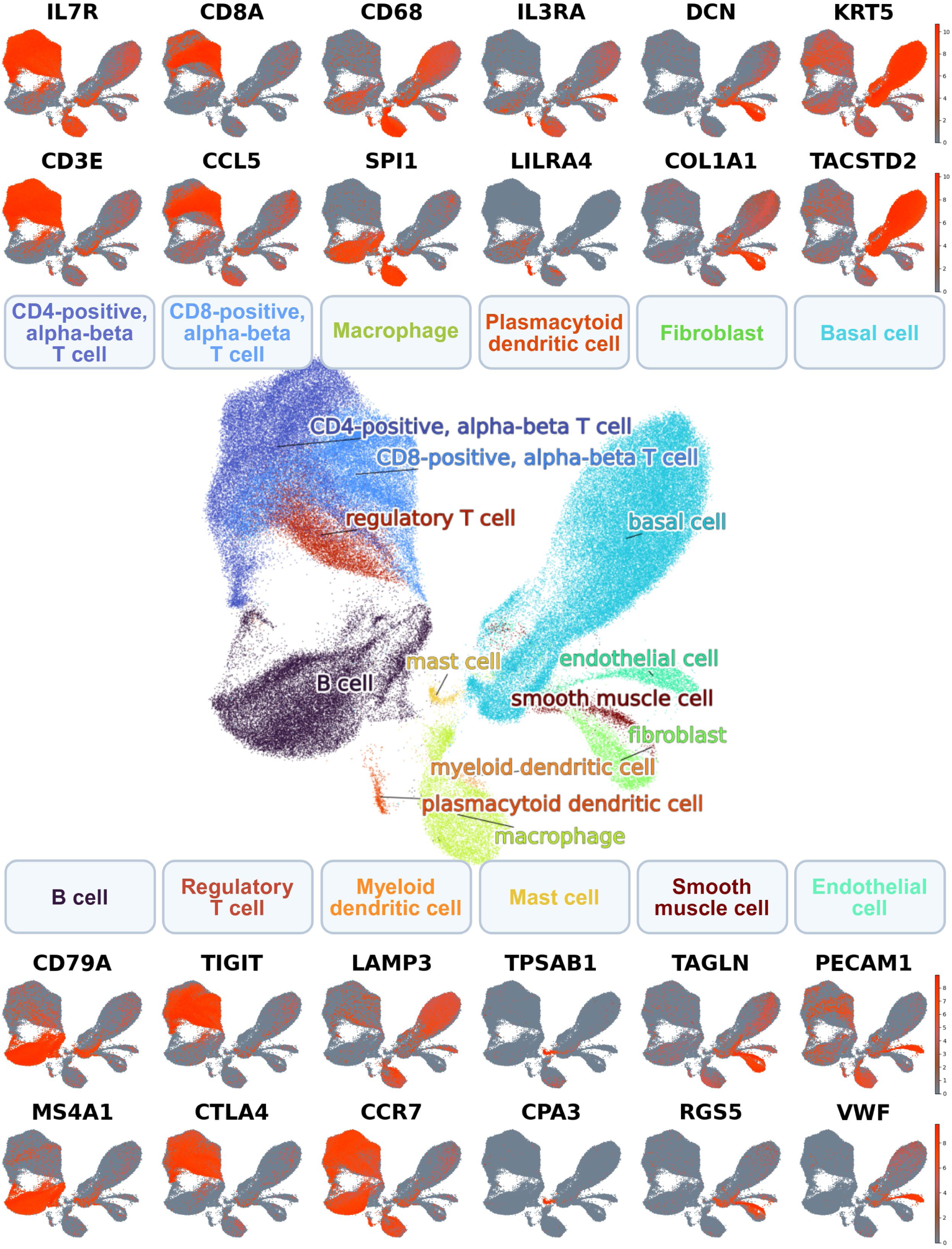
Validation of annotation with known marker genes. UMAPs show the expression of known marker genes that are largely enriched in the annotated cluster. All cell types expressed markers that have been previously reported to characterize these cell types, supporting their annotation. CD4+ T cells: *IL7R*^56^ and *CD3E*^57^. CD8+ T cells: *CD8A*^58^ and *CCL5*^59^. Macrophages: *CDC8*^60^ and *SPI1*^61^. Plasmacytoid dendritic cells: *IL3RA* (CD123)^62^ and *LILRA4* (ILT7)^63^. Fibroblasts: *DCN* and *COL1A1*^64^. Basal cells: *KRT5*^65^ and *TACSTD2*^66^. B cells: *CD7SA* and *MS4A1*^67^. Regulatory T cells: *TIGIT* and *CTLA4* (CD152)^68^. Myeloid dendritic cells: *LAMP3* and *CCR7*^69^. Mast cells: *TPSAB1* and *CPA3*^70^. Smooth muscle cells: *TAGLN* (SM22a)^71^ and *RGS5*^72^. Endothelial cells: *PECAM1* (CD31) and *VWF*^73^.

**Supplemental Figure 4.**
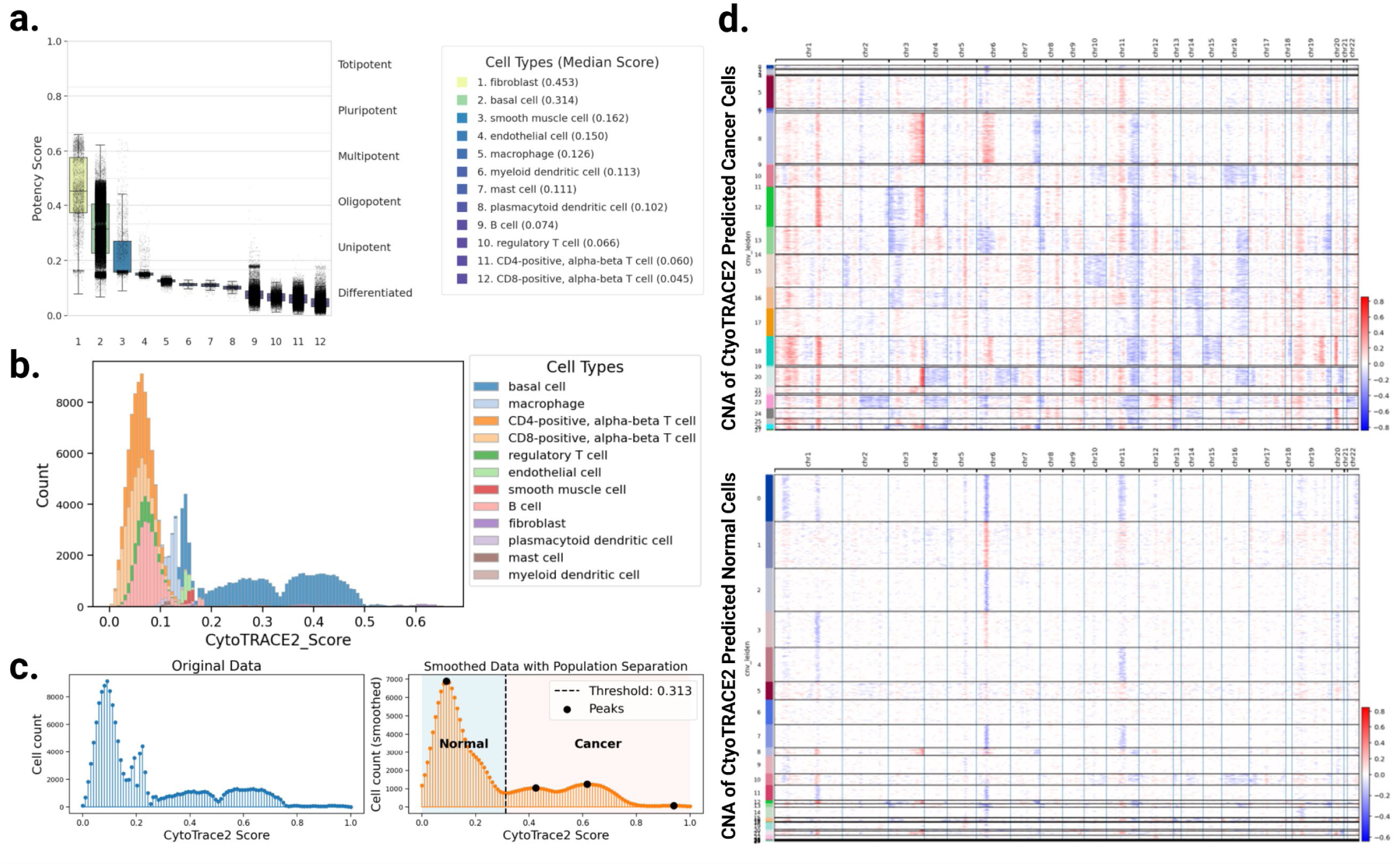
Functional analysis of stemness phenotypes and DNA damage repair in HNSCC in a subpopulation of basal epithelial cells. **(a)** Box and whisker plot of gene expression similarity to stem cells (Potency Score) produced by CytoTRACE 2^26^. We see that for the majority of immune cell types there is one clear population of scores, but for some somatic cell types (basal epithelial cells and fibroblasts) we see two populations of cells with either low or elevated scores. This approach identifies somatic cells in either a normal differentiated cell state or a tumor state with an elevated stem cell transcriptional phenotype. **(b)** Histogram of potency scores stacked by cell type. The histogram shows that the right-sided tail of the potency distribution overlaps with some normal somatic cells near 0.2, above which the potency scores are enriched for basal epithelial cells. **(c)** Original and smoothed histograms with valley detection at the low point between the first two potency peaks. The threshold for normal versus cancer cells was set to distinguish populations for CNA analysis using inferCNV^27^. **(d)** CNA genome tracks showing gain (red) or loss (blue) of predicted CNA alterations, which are hierarchically clustered. We see an overall increase in CNA in our predicted cancer cells, supporting that these cells have higher chromosome instability.

**Supplemental Figure 5.**
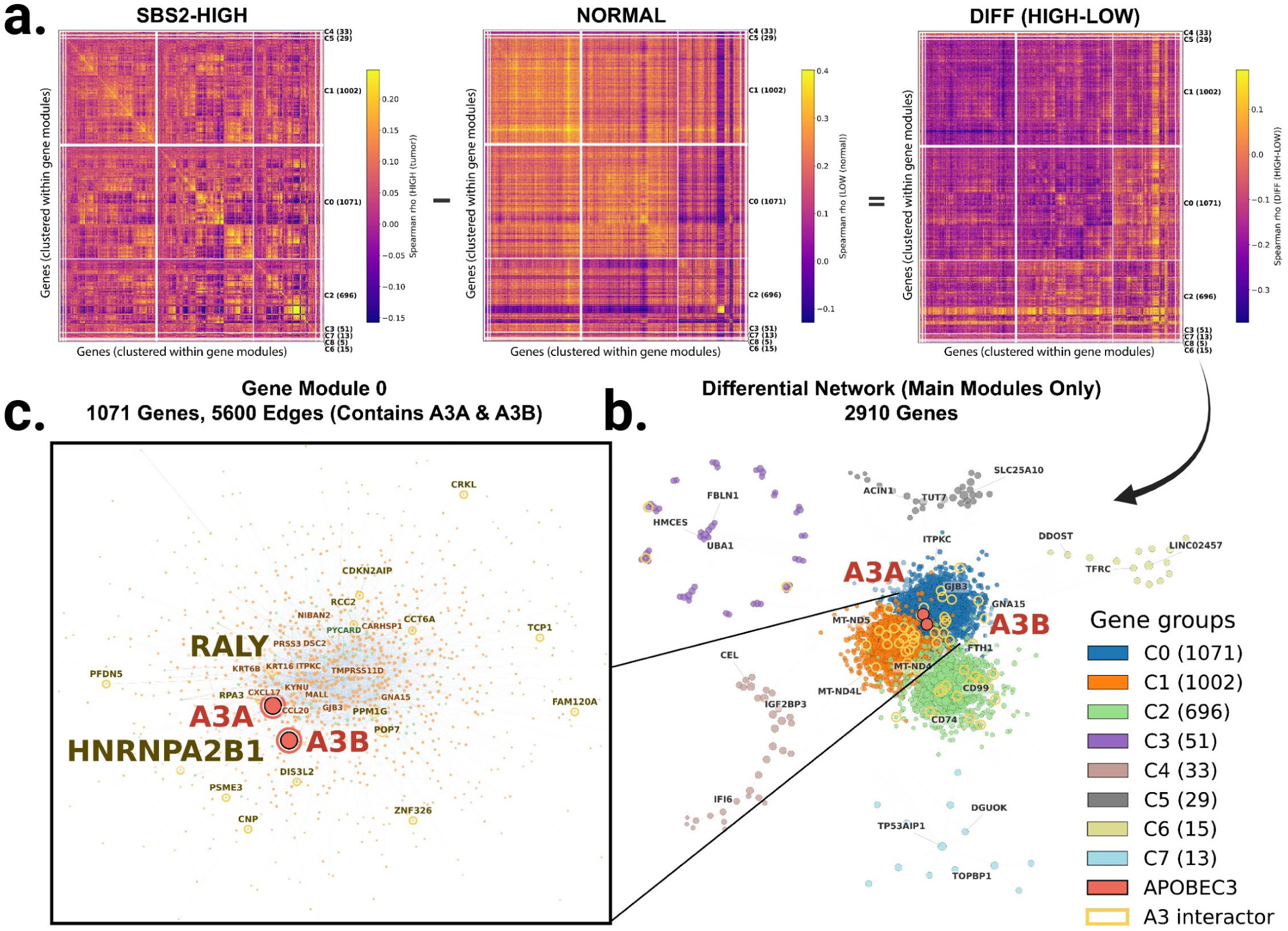
Differential co-expression networks in SBS2-HIGH versus NORMAL basal epithelial cell populations reveal gene communities associated with SBS2 mutagenic activity. (a) Spearman co-expression correlation matrices computed within the SBS2-HIGH group (left) and NORMAL group (center), and the resulting differential (DIFF) matrix (HIGH minus NORMAL; right). The DIFF matrix isolates gene pairs whose co-expression is specifically rewired between SBS2-HIGH tumor and NORMAL tissue conditions. (b) The arrow indicates the global network derived from the DIFF matrix, with main communities labeled. Satellite communities with fewer than 5 genes were excluded from the visualization for clarity. (c) Inset showing Community 0 (1071 genes), the community containing *A3A* and *A3B*.

**Supplemental Figure 6.**
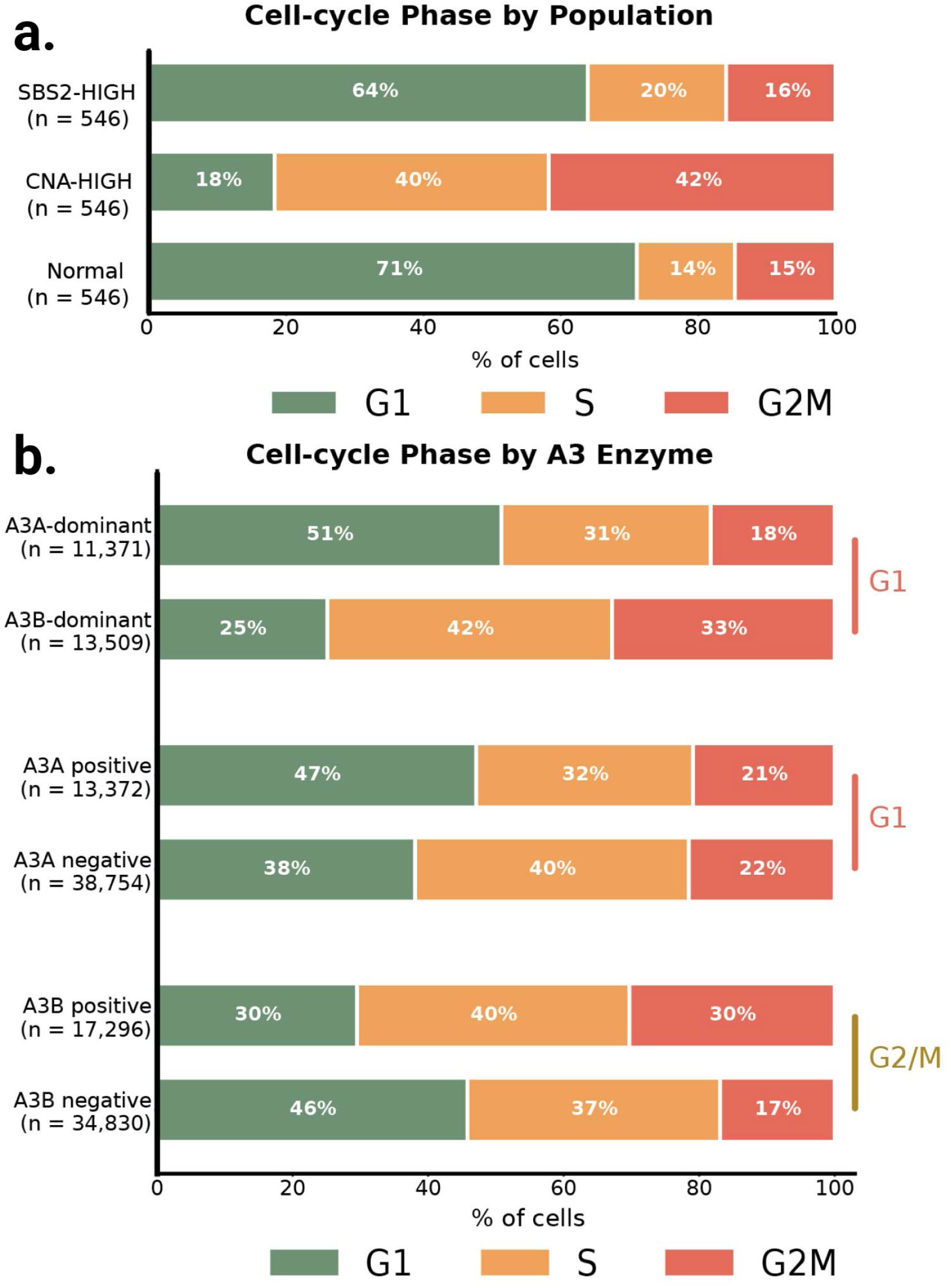
Cell-cycle phase distribution separates the SBS2-HIGH and CNA-HIGH populations and tracks *A3* enzyme dominance and presence/absence in epithelial cells. (a) Relative proportion of cells in each cell-cycle phase for the SBS2-HIGH, CNA-HIGH, and NORMAL epithelial populations. SBS2-HIGH cells are predominantly G1 (63.9%) and resemble NORMAL cells (71.1% G1), while CNA-HIGH cells are predominantly S or G2/M (81.7%, with 41.8% in G2/M). Chi-square = 360.4, Cramér’s V = 0.332. (b) Relative proportion of cells in each cell cycle phase presented in three ways: *A3A*-dominant against *A3B*-dominant cells, defined by *A3A*/(*A3A* + *A3B*) above or below 0.5 among cells expressing either enzyme, *A3A*-positive against *A3A*-negative cells, and *A3B*-positive against *A3B*-negative cells. *A3A*-dominant cells carry 50.8% G1 against 25.1% in *A3B*-dominant cells, a shift of 25.7 percentage points (chi-square = 1814.4, Cramér’s V = 0.270). *A3B*-positive cells are enriched for G2/M relative to *A3B*-negative cells (odds ratio 2.13, 13.4 percentage points) and *A3A*-positive cells for G1 (odds ratio 1.45, 9.0 percentage points).

**Supplemental Figure 7.**
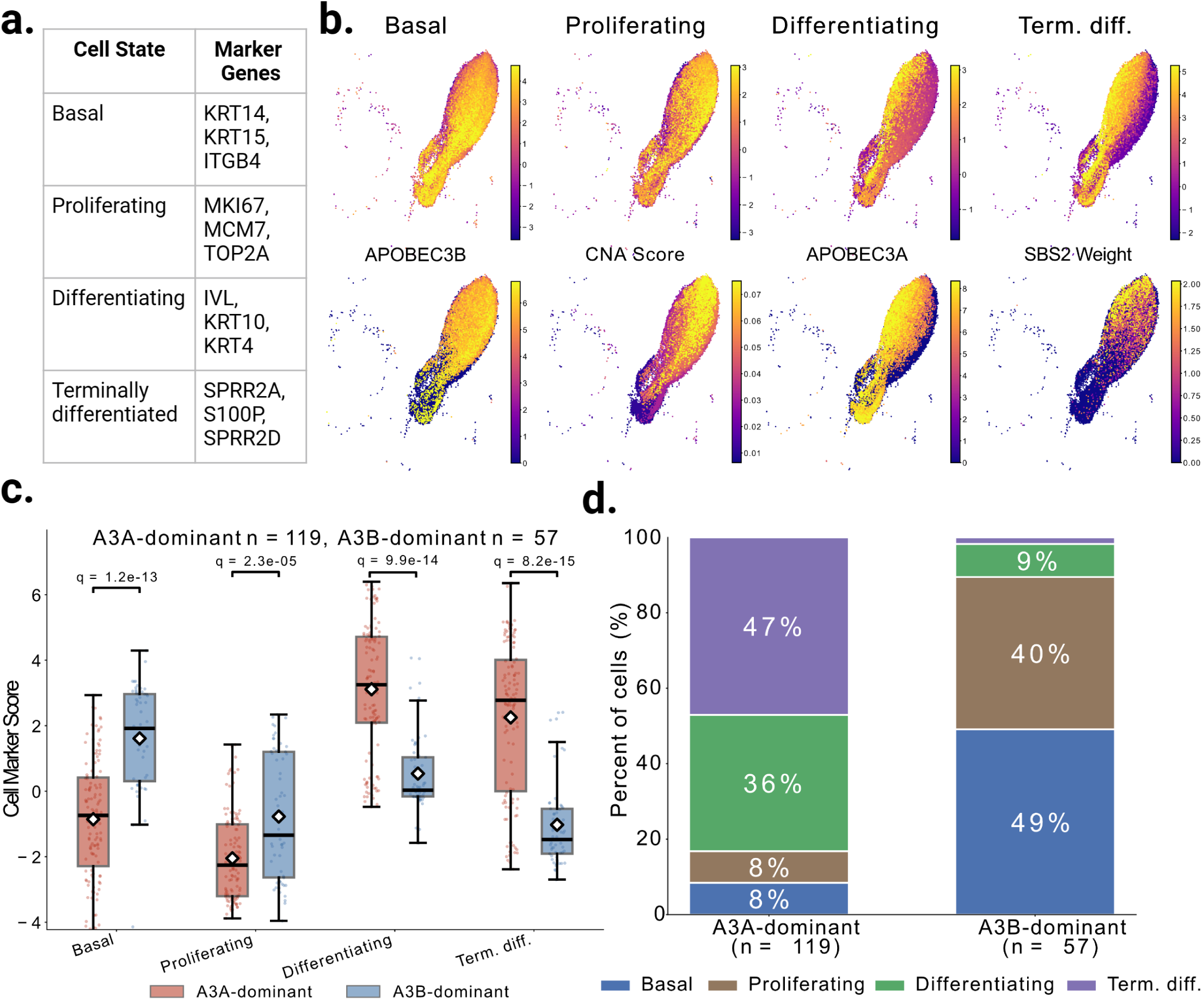
*A3A*– and *A3B*-dominant basal epithelial cells occupy distinct differentiation states in normal-adjacent tissue. **(a)** Marker genes used to define basal, proliferating, differentiating, and terminally differentiated epithelial cell states as previously described^18^. **(b)** Top row, per-cell score for each state across all 52,126 basal epithelial cells, shown on the UMAP embedding. Bottom row, UMAPs displaying *A3B* expression, CNA score, *A3A* expression, and SBS2 weight. **(c)** State scores in normal-adjacent basal epithelial cells by *A3* dominance (*A3A*-dominant n = 119, *A3B*-dominant n = 57). Boxes show the median and interquartile range and white diamonds the mean. Brackets carry Benjamini-Hochberg q-values from two-sided Mann-Whitney U tests corrected across the four states. **(d)** Percentage of the same cells assigned to each state. *A3A*-dominant normal-adjacent epithelial cells were predominantly differentiating or terminally differentiated, whereas *A3B*-dominant normal-adjacent epithelial cells were predominantly basal or proliferating.

**Supplemental Figure 8.**
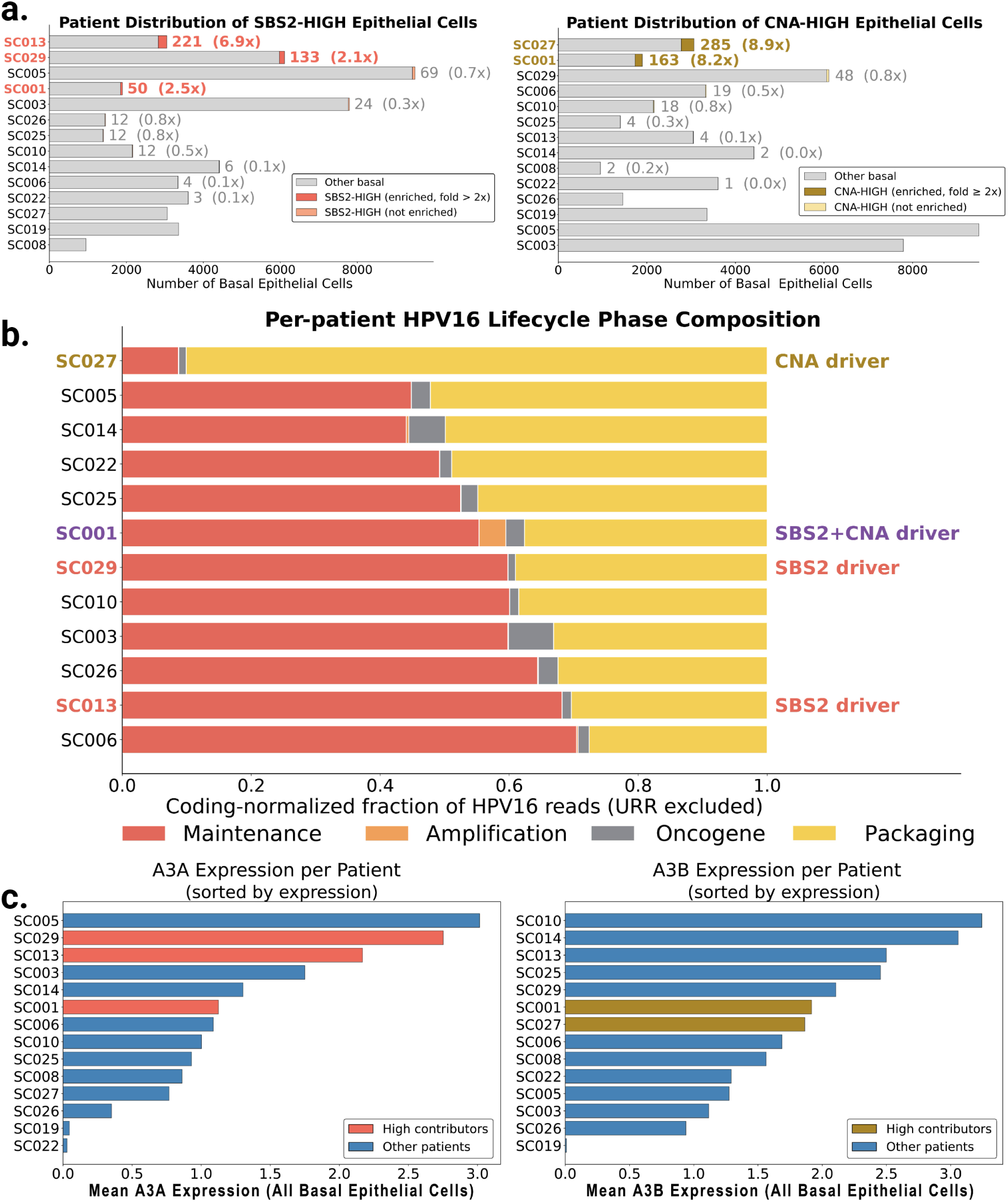
The SBS2-HIGH and CNA-HIGH populations are drawn disproportionately from a subset of patients. **(a)** Cells contributed by each patient to the SBS2-HIGH and CNA-HIGH populations, sorted by contribution. High contributor patients are highlighted. Three patients account for 74.0% of SBS2-HIGH cells (SC013 6.9-fold, SC029 2.1-fold, SC001 2.5-fold; chi-square = 1481.4, df = 13, p = 4.46 x 10^−309^) and two for 82.1% of CNA-HIGH cells (SC027 8.9-fold, SC001 8.2-fold; chi-square = 3419.8, df = 13, p < 1 x 10^−300^). SC001 is the only patient contributing to both. **(b)** Proportion of each patient’s HPV16 coding reads in each functional viral gene group: maintenance (*E1*, *E2*), amplification (*E4*, *E5*), oncogene (*EC*, *E7*), and packaging (*L1*, *L2*). Reads from the upstream regulatory region are excluded, so the bars describe how viral output is distributed rather than how much virus is present. Counts are pooled across each patient’s HPV16-positive cells, and patients with fewer than ten gated cells are unassigned. **(c)** Mean log-normalized *A3A* and *A3B* expression per patient, with high contributors highlighted. The high contributors are not the highest expressors of either enzyme.

**Supplemental Figure 9.**
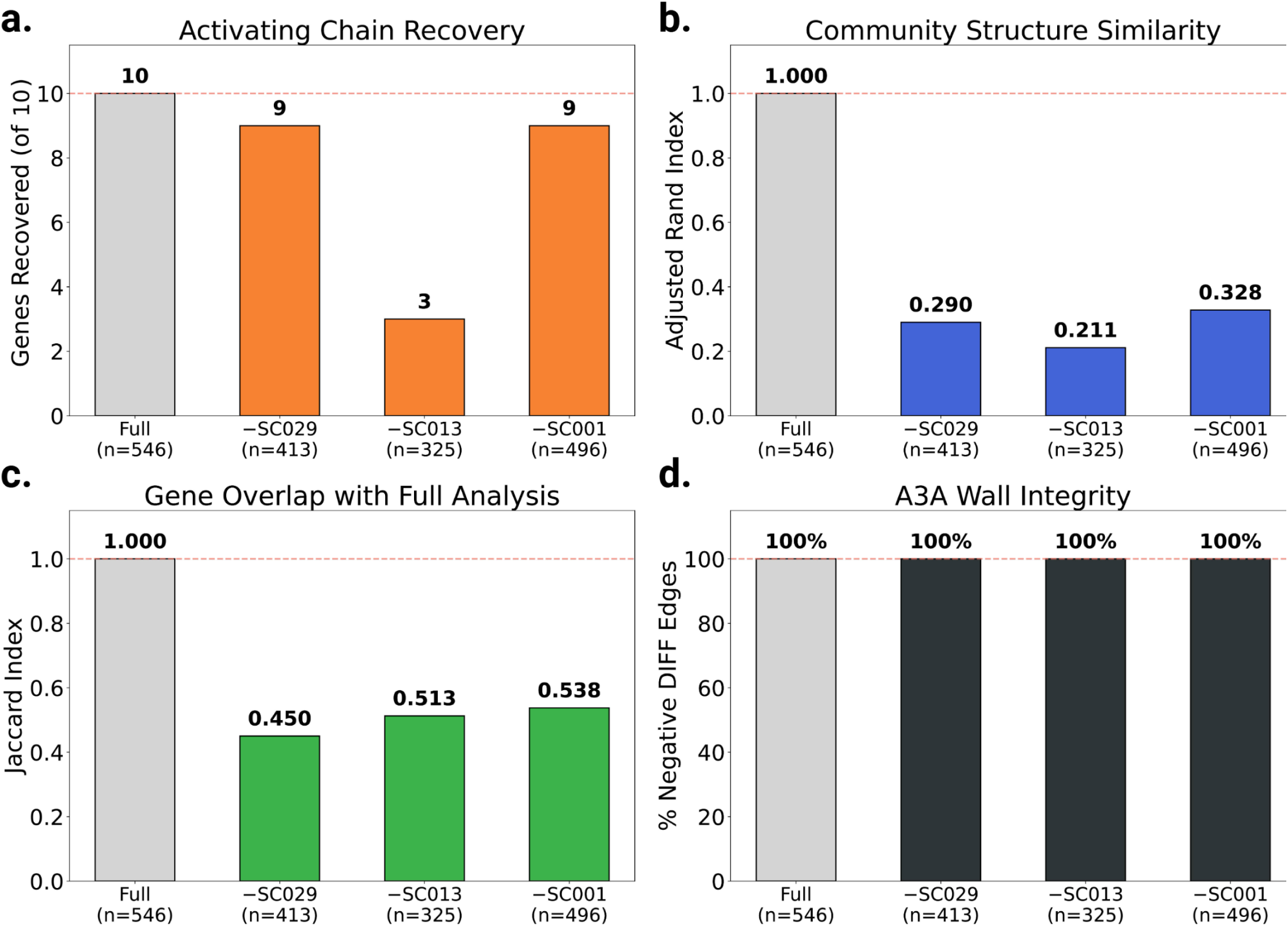
Leave-one-patient-out (LOPO) network sensitivity analysis. Each panel shows the full SBS2-HIGH versus NORMAL network result (leftmost bar) followed by results after removing each high-contributing patient (SC029, SC013, SC001) from the SBS2-HIGH group and re-running the complete network pipeline. **(a)** Activating chain gene recovery (out of 10 genes). Removing SC029 or SC001 retains 9 of 10; removing SC013, which supplies 221 of the 546 SBS2-HIGH cells, retains 3 of 10 (*RALY*, *LCN2*, and *RRAD*). *SMOX* is absent from all three reconstructions. **(b)** Community structure similarity (adjusted Rand index) between the LOPO network and the full analysis ranges from 0.21 to 0.33, indicating community boundaries shift substantially when cells are removed. **(c)** Gene overlap (Jaccard index) between the LOPO network and the full analysis ranged from 0.45 to 0.54. **(d)** A3 wall integrity, measured as the percentage of negative DIFF edges connecting *A3A* and *A3B* to their neighbors. The wall remains 100% intact across all three LOPO configurations, confirming that the pulsatile A3 co-expression signature is not driven by any single patient.

**Supplemental Figure 10.**
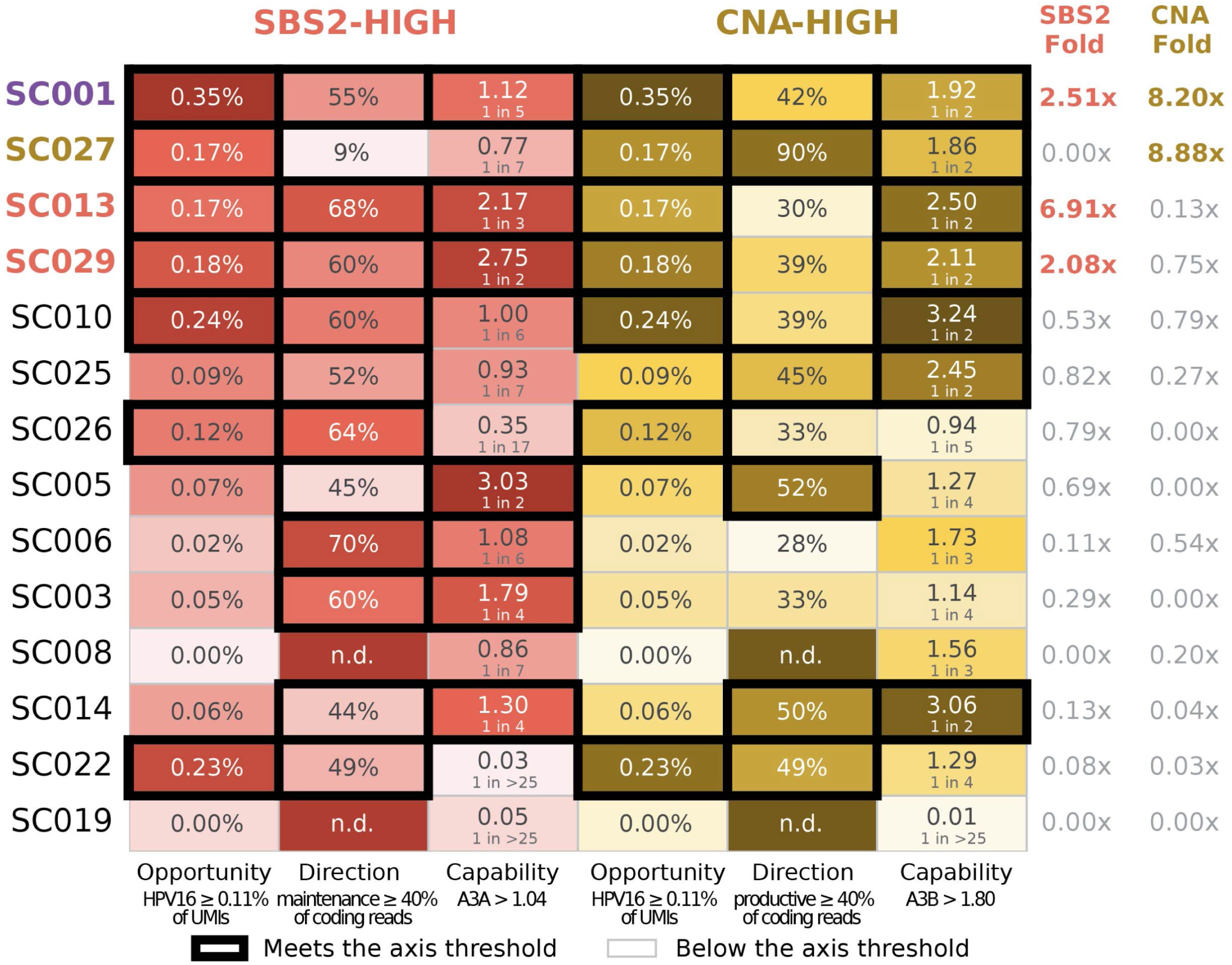
Contribution to each mutagenic fate depends on viral load, life cycle direction, and prevalent expression of the matching A3 enzyme together. Heatmap of per-patient status along the axes that predict contribution. Predictors include HPV16 load as a fraction of epithelial-compartment transcripts, life cycle direction as the fraction of viral coding reads in the stage matching each fate, and *A3* penetrance as the fraction of epithelial cells expressing the matching enzyme. High contributors uniformly meet these qualifications: HPV16 at 0.1% or more of epithelial-compartment transcripts, 40% or more of coding reads in the matching stage, and the matching enzyme expressed in roughly one in five epithelial cells for *A3A* or one in two for *A3B*. All three conditions hold together for every high contributor and for no other patient (Fisher exact p = 0.0027 for SBS2-HIGH, p = 0.011 for CNA-HIGH). Cells meeting the threshold for high contribution carry a thick dark border. No axis is sufficient alone. Examples of this include: patients SC013 and SC027 carry near-identical viral load and go to opposite fates. Patient SC005 carries the highest cohort *A3A* but carries only a third of the average of virus observed in high contributors and contributes to neither. Patient SC010 carries abundant maintenance-stage virus with *A3A* at the cohort median and also contributes to neither.

## Notes

### Competing Interest Statement

The authors have declared no competing interest.

### Summary of Updates

We have performed additional analysis on the normal-adjacent epithelial cells and added a new supplemental figure 7, as well as providing additional edits to wording and citations listed in the text.

## References

1. Sheehy, A.M., Gaddis, N.C., Choi, J.D., and Malim, M.H. (2002). Isolation of a human gene that inhibits HIV-1 infection and is suppressed by the viral Vif protein. Nature 418, 646–650. 10.1038/NATURE00939.

2. Harris, R.S., Bishop, K.N., Sheehy, A.M., Craig, H.M., Petersen-Mahrt, S.K., Watt, I.N., Neuberger, M.S., and Malim, M.H. (2003). DNA Deamination Mediates Innate Immunity to Retroviral Infection. Cell 113, 803–809. 10.1016/S0092-8674(03)00423-9.

3. Mangeat, B., Turelli, P., Caron, G., Friedli, M., Perrin, L., and Trono, D. (2003). Broad antiretroviral defence by human APOBEC3G through lethal editing of nascent reverse transcripts. Nature 2003 424:6944 424, 99–103. 10.1038/nature01709.

4. Alexandrov, L.B., Nik-Zainal, S., Wedge, D.C., Campbell, P.J., and Stratton, M.R. (2013). Deciphering Signatures of Mutational Processes Operative in Human Cancer. Cell Rep. 3, 246. 10.1016/J.CELREP.2012.12.008.

5. Alexandrov, L.B., Kim, J., Haradhvala, N.J., Huang, M.N., Tian Ng, A.W., Wu, Y., Boot, A., Covington, K.R., Gordenin, D.A., Bergstrom, E.N., et al. (2020). The repertoire of mutational signatures in human cancer. Nature 2020 578:7793 578, 94–101. 10.1038/s41586-020-1943-3.

6. Chan, K., Roberts, S.A., Klimczak, L.J., Sterling, J.F., Saini, N., Malc, E.P., Kim, J., Kwiatkowski, D.J., Fargo, D.C., Mieczkowski, P.A., et al. (2015). An APOBEC3A hypermutation signature is distinguishable from the signature of background mutagenesis by APOBEC3B in human cancers. Nat. Genet. 47, 1067–1072. 10.1038/NG.3378.

7. Roberts, S.A., and Gordenin, D.A. (2014). Hypermutation in human cancer genomes: footprints and mechanisms. Nat. Rev. Cancer 14, 786–800. 10.1038/NRC3816.

8. Petljak, M., Green, A.M., Maciejowski, J., and Weitzman, M.D. (2022). Addressing the benefits of inhibiting APOBEC3-dependent mutagenesis in cancer. Nat. Genet. 54, 1599. 10.1038/S41588-022-01196-8.

9. Warren, C.J., Xu, T., Guo, K., Griffin, L.M., Westrich, J.A., Lee, D., Lambert, P.F., Santiago, M.L., and Pyeon, D. (2015). APOBEC3A Functions as a Restriction Factor of Human Papillomavirus. J. Virol. 89, 688–702. 10.1128/JVI.02383-14.

10. Vieira, V.C., Leonard, B., White, E.A., Starrett, G.J., Temiz, N.A., Lorenz, L.D., Lee, D., Soares, M.A., Lambert, P.F., Howley, P.M., et al. (2014). Human papillomavirus E6 triggers upregulation of the antiviral and cancer genomic DNA deaminase APOBEC3B. mBio 5. 10.1128/MBIO.02234-14.

11. Westrich, J.A., Warren, C.J., Klausner, M.J., Guo, K., Liu, C.-W., Santiago, M.L., and Pyeon, D. (2018). Human Papillomavirus 16 E7 Stabilizes APOBEC3A Protein by Inhibiting Cullin 2-Dependent Protein Degradation. J. Virol. 92, 1318–1335. 10.1128/JVI.01318-17.

12. Koning, F.A., Newman, E.N.C., Kim, E.-Y., Kunstman, K.J., Wolinsky, S.M., and Malim, M.H. (2009). Defining APOBEC3 Expression Patterns in Human Tissues and Hematopoietic Cell Subsets. J. Virol. 83, 9474. 10.1128/JVI.01089-09.

13. Bedard, M.C., Chihanga, T., Carlile, A., Jackson, R., Brusadelli, M.G., Lee, D., VonHandorf, A., Rochman, M., Dexheimer, P.J., Chalmers, J., et al. (2023). Single cell transcriptomic analysis of HPV16-infected epithelium identifies a keratinocyte subpopulation implicated in cancer. Nat. Commun. 14, 1975. 10.1038/S41467-023-37377-0.

14. Li, Y., Wang, C., Ma, A., Rani, A.Q., Luo, M., Li, J., Liu, X., and Ma, Q. (2023). Identification of HPV oncogene and host cell differentiation associated cellular heterogeneity in cervical cancer via single-cell transcriptomic analysis. J. Med. Virol. 95. 10.1002/JMV.29060.

15. Puram, S. V., Mints, M., Pal, A., Qi, Z., Reeb, A., Gelev, K., Barrett, T.F., Gerndt, S., Liu, P., Parikh, A.S., et al. (2023). Cellular states are coupled to genomic and viral heterogeneity in HPV-related oropharyngeal carcinoma. Nature Genetics 2023 55:4 55, 640–650. 10.1038/s41588-023-01357-3.

16. Choi, J.H., Lee, B.S., Jang, J.Y., Lee, Y.S., Kim, H.J., Roh, J., Shin, Y.S., Woo, H.G., and Kim, C.H. (2023). Single-cell transcriptome profiling of the stepwise progression of head and neck cancer. Nature Communications 2023 14:1 14, 1055-. 10.1038/s41467-023-36691-x.

17. Kürten, C.H.L., Kulkarni, A., Cillo, A.R., Santos, P.M., Roble, A.K., Onkar, S., Reeder, C., Lang, S., Chen, X., Duvvuri, U., et al. (2021). Investigating immune and non-immune cell interactions in head and neck tumors by single-cell RNA sequencing. Nat. Commun. 12, 7338. 10.1038/S41467-021-27619-4.

18. Smith, N.J., Reddin, I., Policelli, P., Oh, S., Zainal, N., Howes, E., Jenkins, B., Tracy, I., Edmond, M., Sharpe, B., et al. (2025). Differentiation signals induce APOBEC3A expression via GRHL3 in squamous epithelia and squamous cell carcinoma. EMBO J. 44, 1–29. 10.1038/S44318-024-00298-9.

19. Chu, Y., Dai, E., Li, Y., Han, G., Pei, G., Ingram, D.R., Thakkar, K., Qin, J.J., Dang, M., Le, X., et al. (2023). Pan-cancer T cell atlas links a cellular stress response state to immunotherapy resistance. Nat. Med. 29, 1550–1562. 10.1038/S41591-023-02371-Y.

20. Moody, C.A., and Laimins, L.A. (2009). Human Papillomaviruses Activate the ATM DNA Damage Pathway for Viral Genome Amplification upon Differentiation. PLoS Pathog. 5, e1000605. 10.1371/JOURNAL.PPAT.1000605.

21. Roman, B.R., and Aragones, A. (2021). Epidemiology and incidence of HPV-related cancers of the head and neck. J. Surg. Oncol. 124, 920. 10.1002/JSO.26687.

22. Chaturvedi, A.K., Engels, E.A., Pfeiffer, R.M., Hernandez, B.Y., Xiao, W., Kim, E., Jiang, B., Goodman, M.T., Sibug-Saber, M., Cozen, W., et al. (2011). Human Papillomavirus and Rising Oropharyngeal Cancer Incidence in the United States. Journal of Clinical Oncology 29, 4294– 4301. 10.1200/JCO.2011.36.4596.

23. Ergen, C., Xing, G., Xu, C., Kim, M., Jayasuriya, M., McGeever, E., Oliveira Pisco, A., Streets, A., and Yosef, N. (2024). Consensus prediction of cell type labels in single-cell data with popV. Nature Genetics 2024 56:12 56, 2731–2738. 10.1038/s41588-024-01993-3.

24. Warren, C.J., Westrich, J.A., Van Doorslaer, K., and Pyeon, D. (2017). Roles of APOBEC3A and APOBEC3B in Human Papillomavirus Infection and Disease Progression. Viruses 2017, Vol. 9, Page 233 *9*, 233. 10.3390/V9080233.

25. Muyas, F., Sauer, C.M., Valle-Inclán, J.E., Li, R., Rahbari, R., Mitchell, T.J., Hormoz, S., and Cortés-Ciriano, I. (2023). De novo detection of somatic mutations in high-throughput single-cell profiling data sets. Nat. Biotechnol. 42, 758. 10.1038/S41587-023-01863-Z.

26. Kang, M., Gulati, G.S., Brown, E.L., Qi, Z., Avagyan, S., Armenteros, J.J.A., Gleyzer, R., Zhang, W., Steen, C.B., D’Silva, J.P., et al. (2025). Improved reconstruction of single-cell developmental potential with CytoTRACE 2. Nature Methods 2025 22:11 22, 2258–2263. 10.1038/s41592-025-02857-2.

27. Patel, A.P., Tirosh, I., Trombetta, J.J., Shalek, A.K., Gillespie, S.M., Wakimoto, H., Cahill, D.P., Nahed, B. V., Curry, W.T., Martuza, R.L., et al. (2014). Single-cell RNA-seq highlights intratumoral heterogeneity in primary glioblastoma. Science 344, 1396. 10.1126/SCIENCE.1254257.

28. Petljak, M., Alexandrov, L.B., Brammeld, J.S., Price, S., Wedge, D.C., Grossmann, S., Dawson, K.J., Ju, Y.S., Iorio, F., Tubio, J.M.C., et al. (2019). Characterizing Mutational Signatures in Human Cancer Cell Lines Reveals Episodic APOBEC Mutagenesis. Cell 176, 1282. 10.1016/J.CELL.2019.02.012.

29. Kanehisa, M., and Goto, S. (2000). KEGG: Kyoto Encyclopedia of Genes and Genomes. Nucleic Acids Res. 28, 27. 10.1093/NAR/28.1.27.

30. McCann, J.L., Cristini, A., Law, E.K., Lee, S.Y., Tellier, M., Carpenter, M.A., Beghè, C., Kim, J.J., Sanchez, A., Jarvis, M.C., et al. (2023). APOBEC3B regulates R-loops and promotes transcription-associated mutagenesis in cancer. Nature Genetics 2023 55:10 55, 1721–1734. 10.1038/s41588-023-01504-w.

31. Jang, G.M., Sudarsan, A.K.A., Shayeganmehr, A., Munhoz, E.P., Lao, R., Gaba, A., Rodríguez, M.G., Love, R.P., Polacco, B.J., Zhou, Y., et al. (2024). Protein Interaction Map of APOBEC3 Enzyme Family Reveals Deamination-Independent Role in Cellular Function. Molecular and Cellular Proteomics 23. 10.1016/j.mcpro.2024.100755.

32. Müller-Bötticher, N., Sahay, S., Eils, R., and Ishaque, N. (2025). SpatialLeiden: spatially aware Leiden clustering. Genome Biol. 26, 24. 10.1186/S13059-025-03489-7.

33. Schichl, K., and Doorbar, J. (2025). Regulation and Deregulation of Viral Gene Expression During High-Risk HPV Infection. Viruses 17, 937. 10.3390/V17070937.

34. Doorbar, J., Quint, W., Banks, L., Bravo, I.G., Stoler, M., Broker, T.R., and Stanley, M.A. (2012). The biology and life-cycle of human papillomaviruses. Vaccine 30. 10.1016/j.vaccine.2012.06.083.

35. Hong, S., Cheng, S., Iovane, A., and Laimins, L.A. (2015). STAT-5 Regulates Transcription of the Topoisomerase IIβ-Binding Protein 1 (TopBP1) Gene To Activate the ATR Pathway and Promote Human Papillomavirus Replication. mBio 6, e02006–15. 10.1128/MBIO.02006-15.

36. Moody, C.A., Fradet-Turcotte, A., Archambault, J., and Laimins, L.A. (2007). Human papillomaviruses activate caspases upon epithelial differentiation to induce viral genome amplification. Proc. Natl. Acad. Sci. U. S. A. 104, 19541–19546. 10.1073/PNAS.0707947104.

37. Wu, S.Y., Lai, H.T., Sanjib Banerjee, N., Ma, Z., Santana, J.F., Wei, S., Liu, X., Zhang, M., Zhan, J., Chen, H., et al. (2024). IDR-targeting compounds suppress HPV genome replication via disruption of phospho-BRD4 association with DNA damage response factors. Mol. Cell 84, 202–220.e15. 10.1016/J.MOLCEL.2023.11.022.

38. Hirabayashi, S., Shirakawa, K., Horisawa, Y., Matsumoto, T., Matsui, H., Yamazaki, H., Sarca, A.D., Kazuma, Y., Nomura, R., Konishi, Y., et al. (2021). APOBEC3B is preferentially expressed at the G2/M phase of cell cycle. Biochem. Biophys. Res. Commun. 546, 178–184. 10.1016/J.BBRC.2021.02.008.

39. Wu, C.H., Zhou, X., and Chen, M. (2025). Exploring and mitigating shortcomings in single-cell differential expression analysis with a new statistical paradigm. Genome Biol. 26, 58. 10.1186/S13059-025-03525-6.

40. Xie, N., Shen, G., Gao, W., Huang, Z., Huang, C., and Fu, L. (2023). Neoantigens: promising targets for cancer therapy. Signal Transduction and Targeted Therapy 2022 8:1 8, 9-. 10.1038/s41392-022-01270-x.

41. Raghavan, M., Zaitoua, A.J., and Kaur, A. (2020). Variations in MHC class I antigen presentation and immunopeptidome selection pathways. F1000Res. 9, F1000 Faculty Rev-1177. 10.12688/F1000RESEARCH.26935.1.

42. Sethna, Z., Guasp, P., Reiche, C., Milighetti, M., Ceglia, N., Patterson, E., Lihm, J., Payne, G., Lyudovyk, O., Rojas, L.A., et al. (2025). RNA neoantigen vaccines prime long-lived CD8+ T cells in pancreatic cancer. Nature 2025 639:8056 639, 1042–1051. 10.1038/s41586-024-08508-4.

43. Wu, B., Zhang, B., Li, B., Wu, H., and Jiang, M. (2024). Cold and hot tumors: from molecular mechanisms to targeted therapy. Signal Transduction and Targeted Therapy 2024 9:1 9, 274-. 10.1038/s41392-024-01979-x.

44. Zheng, G.X.Y., Terry, J.M., Belgrader, P., Ryvkin, P., Bent, Z.W., Wilson, R., Ziraldo, S.B., Wheeler, T.D., McDermott, G.P., Zhu, J., et al. (2017). Massively parallel digital transcriptional profiling of single cells. Nature Communications 2017 8:1 8, 14049-. 10.1038/ncomms14049.

45. Wolock, S.L., Lopez, R., and Klein, A.M. (2019). Scrublet: Computational Identification of Cell Doublets in Single-Cell Transcriptomic Data. Cell Syst. 8, 281–291.e9. 10.1016/j.cels.2018.11.005.

46. Jones, R.C., Karkanias, J., Krasnow, M.A., Pisco, A.O., Quake, S.R., Salzman, J., Yosef, N., Bulthaup, B., Brown, P., Harper, W., et al. (2022). The Tabula Sapiens: A multiple-organ, single-cell transcriptomic atlas of humans. Science (1979). 376. 10.1126/SCIENCE.ABL4896.

47. Wolf, F.A., Angerer, P., and Theis, F.J. (2018). SCANPY: large-scale single-cell gene expression data analysis. Genome Biol. 19. 10.1186/S13059-017-1382-0.

48. Kuleshov, M. V., Jones, M.R., Rouillard, A.D., Fernandez, N.F., Duan, Q., Wang, Z., Koplev, S., Jenkins, S.L., Jagodnik, K.M., Lachmann, A., et al. (2016). Enrichr: a comprehensive gene set enrichment analysis web server 2016 update. Nucleic Acids Res. 44, W90–W97. 10.1093/NAR/GKW377.

49. Fang, Z., Liu, X., and Peltz, G. (2023). GSEApy: a comprehensive package for performing gene set enrichment analysis in Python. Bioinformatics 39. 10.1093/BIOINFORMATICS/BTAC757.

50. Wood, D.E., Lu, J., and Langmead, B. (2019). Improved metagenomic analysis with Kraken 2. Genome Biology 2019 20:1 20, 257-. 10.1186/S13059-019-1891-0.

51. Li, H. (2018). Minimap2: pairwise alignment for nucleotide sequences. Bioinformatics 34, 3094–3100. 10.1093/BIOINFORMATICS/BTY191.

52. Cingolani, P., Platts, A., Wang, L.L., Coon, M., Nguyen, T., Wang, L., Land, S.J., Lu, X., and Ruden, D.M. (2012). A program for annotating and predicting the effects of single nucleotide polymorphisms, SnpEff: SNPs in the genome of Drosophila melanogaster strain w1118; iso-2; iso-3. Fly (Austin). 6, 80. 10.4161/FLY.19695.

53. Dyer, S.C., Austine-Orimoloye, O., Azov, A.G., Barba, M., Barnes, I., Barrera-Enriquez, V.P., Becker, A., Bennett, R., Beracochea, M., Berry, A., et al. (2025). Ensembl 2025. Nucleic Acids Res. 53, D948–D957. 10.1093/NAR/GKAE1071.

54. O’Donnell, T.J., Rubinsteyn, A., and Laserson, U. (2020). MHCflurry 2.0: Improved Pan-Allele Prediction of MHC Class I-Presented Peptides by Incorporating Antigen Processing. Cell Syst. 11, 42–48.e7. 10.1016/j.cels.2020.06.010.

55. Bui, H.H., Sidney, J., Dinh, K., Southwood, S., Newman, M.J., and Sette, A. (2006). Predicting population coverage of T-cell epitope-based diagnostics and vaccines. BMC Bioinformatics 7. 10.1186/1471-2105-7-153.

56. Azizi, G., Van den Broek, B., Ishikawa, L.L.W., Naziri, H., Yazdani, R., Zhang, G.X., Ciric, B., and Rostami, A. (2024). IL-7Rα on CD4+ T cells is required for their survival and the pathogenesis of experimental autoimmune encephalomyelitis. J. Neuroinflammation 21, 253. 10.1186/s12974-024-03224-2.

57. Wang, X., Shen, X., Chen, S., Liu, H., Hong, N., Zhong, H., Chen, X., and Jin, W. (2022). Reinvestigation of Classic T Cell Subsets and Identification of Novel Cell Subpopulations by Single-Cell RNA Sequencing. The Journal of Immunology 208, 396–406. 10.4049/jimmunol.2100581.

58. Deng, W., Ma, Y., Su, Z., Liu, Y., Liang, P., Huang, C., Liu, X., Shao, J., Zhang, Y., Zhang, K., et al. (2020). Single-cell RNA-sequencing analyses identify heterogeneity of CD8+ T cell subpopulations and novel therapy targets in melanoma. Mol. Ther. Oncolytics 20, 105. 10.1016/j.omto.2020.12.003.

59. Pauken, K.E., Shahid, O., Lagattuta, K.A., Mahuron, K.M., Luber, J.M., Lowe, M.M., Huang, L., Delaney, C., Long, J.M., Fung, M.E., et al. (2021). Single-cell analyses identify circulating anti-tumor CD8 T cells and markers for their enrichment. Journal of Experimental Medicine 218. 10.1084/JEM.20200920.

60. Mould, K.J., Jackson, N.D., Henson, P.M., Seibold, M., and Janssen, W.J. (2019). Single cell RNA sequencing identifies unique inflammatory airspace macrophage subsets. JCI Insight 4. 10.1172/jci.insight.126556.

61. Hume, D.A., Millard, S.M., and Pettit, A.R. (2023). Macrophage heterogeneity in the single-cell era: facts and artifacts. Blood 142, 1339–1347. 10.1182/blood.2023020597.

62. Jegalian, A.G., Facchetti, F., and Jaffe, E.S. (2009). Plasmacytoid Dendritic Cells: Physiologic Roles and Pathologic States. Adv. Anat. Pathol. 16, 392. 10.1097/PAP.0b013e3181bb6bc2.

63. Tavano, B., and Boasso, A. (2014). Effect of Immunoglobin-Like Transcript 7 Cross-Linking on Plasmacytoid Dendritic Cells Differentiation into Antigen-Presenting Cells. PLoS One *9*, e89414. 10.1371/journal.pone.0089414.

64. Lujano Olazaba, O., Farrow, J., and Monkkonen, T. (2024). Fibroblast heterogeneity and functions: insights from single-cell sequencing in wound healing, breast cancer, ovarian cancer and melanoma. Front. Genet. 15, 1304853. 10.3389/fgene.2024.1304853.

65. Huang, H., Tan, K. Sen, Zhou, S., Yuan, T., Liu, J., Ong, H.H., Chen, Q., Gao, J., Xu, M., Zhu, Z., et al. (2020). p63+Krt5+ basal cells are increased in the squamous metaplastic epithelium of patients with radiation-induced chronic Rhinosinusitis. Radiat. Oncol. 15, 222. 10.1186/s13014-020-01656-7.

66. Goldstein, A.S., Lawson, D.A., Cheng, D., Sun, W., Garraway, I.P., and Witte, O.N. (2008). Trop2 identifies a subpopulation of murine and human prostate basal cells with stem cell characteristics. Proc. Natl. Acad. Sci. U. S. A. 105, 20882–20887. 10.1073/pnas.0811411106.

67. Hu, H., Zhan, W., Zhu, H., Hao, B., Yan, T., Zhang, J., Wang, S., and Zhang, T. (2025). Single-Cell Sequence and Machine Learning Identify a CD79A+B Cells-Related Transcriptional Signature for Predicting Clinical Outcomes and Immune Microenvironment in Breast Cancer. Cancer Inform. 24, 11769351251360676. 10.1177/11769351251360675.

68. Wegrzyn, A.S., Kedzierska, A.E., and Obojski, A. (2023). Identification and classification of distinct surface markers of T regulatory cells. Front. Immunol. 13, 1055805. 10.3389/fimmu.2022.1055805.

69. Guimarães, G.R., Maklouf, G.R., Teixeira, C.E., de Oliveira Santos, L., Tessarollo, N.G., de Toledo, N.E., Serain, A.F., de Lanna, C.A., Pretti, M.A., da Cruz, J.G.V., et al. (2024). Single-cell resolution characterization of myeloid-derived cell states with implication in cancer outcome. Nature Communications 2024 15:1 15, 5694-. 10.1038/s41467-024-49916-4.

70. Chantran, Y., and Arock, M. (2025). Hereditary alpha-tryptasemia and monoclonal mast cell disorders. Frontiers in Allergy 6, 1600680. 10.3389/falgy.2025.1600680.

71. Tsuji-Tamura, K., Morino-Koga, S., Suzuki, S., and Ogawa, M. (2021). The canonical smooth muscle cell marker TAGLN is present in endothelial cells and is involved in angiogenesis. J. Cell Sci. 134. 10.1242/JCS.254920.

72. Gao, Y.K., Guo, R.J., Xu, X., Huang, X.F., Song, Y., Zhang, D.D., Chen, N., Wang, X.W., Liang, C.X., Kong, P., et al. (2022). A regulator of G protein signaling 5 marked subpopulation of vascular smooth muscle cells is lost during vascular disease. PLoS One 17, e0265132. 10.1371/journal.pone.0265132.

73. Müller, A.M., Hermanns, M.I., Skrzynski, C., Nesslinger, M., Müller, K.M., and Kirkpatrick, C.J. (2002). Expression of the endothelial markers PECAM-1, vWF, and CD34 in Vivo and in Vitro. Exp. Mol. Pathol. 72, 221–229. 10.1006/exmp.2002.2424.

